# Towards optimised extracellular vesicle proteomics from cerebrospinal fluid

**DOI:** 10.1101/2022.09.27.508990

**Authors:** Petra Kangas, Tuula A. Nyman, Liisa Metsähonkala, Cameron Burns, Robert Tempest, Tim Williams, Jenni Karttunen, Tarja S. Jokinen

## Abstract

The proteomic profile of extracellular vesicles (EVs) from cerebrospinal fluid (CSF) can reveal novel biomarkers for diseases of the brain. Here, we validate an ultrafiltration combined with size-exclusion chromatography (UF-SEC) method for isolation of EVs from canine CSF and probe the effect of starting volume on the EV proteomics profile. First, we performed a literature review of CSF EV articles to define the current state of art, discovering a need for basic characterisation of CSF EVs. Secondly, we isolated EVs from CSF by UF-SEC and characterised the SEC fractions by protein amount, particle count, transmission electron microscopy, and immunoblotting. Data are presented as mean ± standard deviation. Using proteomics, SEC fractions 3-5 were compared and enrichment of EV markers in fraction 3 was detected, whereas fractions 4-5 contained more apolipoproteins. Lastly, we compared starting volumes of pooled CSF (6ml, 3ml, 1ml, and 0.5ml) to evaluate the effect on the proteomic profile. Even with a 0.5ml starting volume, 743±77 or 345±88 proteins were identified depending on whether ‘matches between runs’ was active in MaxQuant. The results confirm that UF-SEC effectively isolates CSF EVs and that EV proteomic analysis can be performed from 0.5ml of canine CSF.

## Introduction

Extracellular vesicle (EV) is an umbrella term for various kinds of cell-released membranous particles. Traditional EV subtypes include exosomes, that are secreted from endosomal pathway, microvesicles from plasma membrane budding, and apoptotic bodies released mainly during cell death (Crescitelli et al., 2020; Théry et al., 2018). Recently, other extracellular nanoparticles, including small (<50nm) non-membranous exomers and supermeres have also been discovered (Zhang et al., 2021, 2019, 2018). Alternative EV subgroups have been classified based on the size and density of the particles since these are properties that can be used to fractionate EVs (Lischnig et al., 2022; Crescitelli et al., 2020).

EVs are a promising tool for biomarker discovery, since EV cargo reflects that of the cells the EVs are secreted from, and it is possible to isolate them from different biofluids. Cerebrospinal fluid (CSF) is a liquid located close to the central nervous system that is secreted mostly by cells in the choroid plexus (Spector et al., 2015; Sturges, 2015; Sakka et al., 2011). An important role of EVs has been demonstrated for the central nervous system, such as regulation of synaptic communication, synaptic strength, and nerve regeneration (Upadhya and Shetty, 2021; Chiasserini et al., 2014). Thus, analysis of EVs secreted into CSF can offer a way to more non-invasively study changes in brain tissue. So far changes in CSF EV cargo have been detected in patients with Alzheimer’s disease and multiple sclerosis (Sandau et al., 2022; Gelibter et al., 2021; Muraoka et al., 2020; Geraci et al., 2018), thus providing proof of concept that CSF derived EVs can be used as a source of biomarkers.

There are various methods available for isolation of EVs, including differential centrifugation, precipitation and size-exclusion chromatography (SEC). Different isolation methods have been described in more detail in a review by Sidhom et al. (Sidhom et al., 2020). The selection of isolation method is a balance between EV yield and purity, which poses challenges for EV biomarker discovery; optimising purity is likely to enhance test specificity, whilst optimising EV yield is necessary to maximise test sensitivity. Ultrafiltration combined with size-exclusion chromatography (UF-SEC) outperforms the differential centrifugation method and/or commercial precipitation-based kits for EV isolation from cell culture supernatant, and urine in terms of purity (Allelein et al., 2021; Karttunen et al., 2021; Sidhom et al., 2020). For CSF EVs, UF-SEC has been reported to increase both yield and purity (Thompson et al., 2018). Density gradient ultracentrifugation produces highly pure EV preparations but is time consuming and requires expensive ultracentrifugation facilities that might not be available in all clinical settings (Ayala-Mar et al., 2019). SEC, on the other hand, is relatively fast and cost-effective, and requires no specialised laboratory equipment (Sidhom et al., 2020). However, it is not capable of separating EVs from similarly sized lipoprotein particles, the latter of which are especially abundant in plasma (Karimi et al., 2018; Simonsen, 2017).

CSF derived EVs have not been widely studied to date, likely because of the invasive collection method and limited volume that can be obtained from individual patients, and because it is difficult to acquire samples from healthy control patients. In a survey performed by International society for Extracellular Vesicles (ISEV) Rigor and Standardization Subcommittee in 2019, only 10% of the respondents had used CSF in their EV studies, which was only a few percentages more compared to the previous survey performed in 2015 (Royo et al., 2020). Also, lipoprotein contamination is a problem with CSF samples. ApoA1 and ApoA2 that are present in CSF are suggested to originate from plasma, and ApoE and ApoJ are suggested to be made within the blood-brain-barrier (Ladu et al., 2000). In addition, small quantities of plasma proteins, such as albumin, are present in CSF (Spector et al., 2015) and can cause contamination.

Canine spontaneous diseases are utilised in translational research and therefore studying canine CSF can be beneficial also for human studies. In dogs, limited amounts of CSF can be collected (recommended 1ml maximum per 5kg body weight), especially from small individuals (Di Terlizzi and Platt, 2009). Hence, more information about the minimum CSF starting volume for isolation of a sufficient amount of EVs is needed. Compared to humans, it is easier to collect high volumes of fresh CSF from deceased animals, which enables methodological studies.

Our overall objective was to validate the UF-SEC method for isolation of CSF derived EVs and to determine the minimum CSF volume required for EV proteomic analysis. We first performed a literature review to investigate the current state of CSF EV research, which revealed a need for basic characterisation of EVs. Secondly, we used the UF-SEC method for EV isolation from canine CSF and used a variety of techniques to confirm the presence of EVs and characterise CSF derived EVs. Finally, we compared the EV proteomes obtained from different volumes of CSF in order to find the minimum volume needed for successful proteomic analysis from CSF EVs. Our results indicate that the UF-SEC protocol can be used to isolate purified EVs from CSF, and that a sufficient amount of EVs can be isolated from 0.5ml of CSF to permit EV proteomic analysis. To our knowledge, this is the first article describing EV isolation from canine CSF.

## Materials and methods

### Literature review

For the literature review, we searched articles about CSF EVs from the following online databases: Web of Science, PubMed, EV Track, Wiley Online Library and Google Scholar using the search term cerebrospinal fluid combined with the terms extracellular vesicles, exosomes, microvesicles, ectosomes and apoptotic bodies. An asterisk was used when necessary to broaden the search results. We only included original research articles published in 2015 or later in the study and the searches were done between April 2021 and August 2022. We reviewed the articles for the starting volume of CSF, for the isolation and characterisation methods, and the number of different characterisation methods used per study. The objectives of the studies were also noted. Articles analysed for the literature review can be found in the **Supplementary file 1**.

### Cerebrospinal fluid collection

Cerebrospinal fluid was collected from dogs euthanised at the Veterinary Teaching Hospital of the University of Helsinki, Finland. The study was approved by the Viikki Campus Research Ethics Committee at the University of Helsinki, Finland. The dogs were euthanised for various reasons unrelated to our study and all the carcasses were donated to teaching and research purposes with written owner consent from the owner.

CSF was collected via a cisternal tap with a sterile puncture within one hour of euthanasia. As much CSF as possible was drained from each dog into 2ml plain tubes. Samples were visually assessed to be clear of any visible red blood cell contamination. Within 30 minutes of collection, total nucleated cell count and total protein concentration were determined. The remainder of the sample was centrifuged at 4000 x g for 10 minutes at room temperature to pellet any cellular material. The supernatant was transferred into a new tube, from where it was divided into Protein LoBind tubes (#022431081, Eppendorf AG, Hamburg, Germany), snap-frozen in liquid nitrogen and stored at –80°C until further use.

### Extracellular vesicle isolation

Prior to EV isolation, CSF aliquots were thawed on ice, pooled and 1/100 volume of protease inhibitor (#P8340, Sigma-Aldrich, St. Louis, Missouri, United States) was added. The pool was centrifuged at 4000 x g for 20 minutes at 4°C to remove any remaining debris. For method set-up, two 7ml pools of CSF were used. For proteomic analysis, three separate pools of 12ml were divided into four volumes: 6ml, 3ml, 1ml, and 0.5ml, which were then processed individually.

For the UF-SEC isolation of EVs, the samples were first concentrated by centrifugation at 4,000 x g at 21°C using Amicon® Ultra-4 Centrifugal Filter Unit with 100kDa molecular weight cutoff (UFC810024, Merck Millipore, Burlington, Massachusetts, United States). After sample concentration, the final volume of each sample was adjusted to 130μl with PBS. SEC was performed with qEV single 70nm columns (Izon, Christchurch, New Zealand), according to manufacturer’s protocol. Briefly, the column was rinsed with a minimum of 10ml of PBS and 130μl of the concentrated sample was loaded into the column. PBS was added as needed and nine 500μl fractions were collected.

Before the proteomics comparisons, an aliquot of 50μl was taken from each fraction for nanoparticle tracking analysis (NTA) and bicinchoninic acid assay (BCA) analysis and the rest of the fraction was stored at -80 °C prior to proteomic analysis.

### Characterisation

#### BCA Protein Assay

The protein concentration of each fraction was measured using the Pierce BCA Protein Assay Kit (#23227, Thermo Scientific™, Waltham, Massachusetts, United States) according to the manufacturer’s instructions. Briefly, two sets of standards and 25μl of each sample were mixed with 200μl of working reagent on a well plate. The plate was incubated at 37°C for 30 minutes and read at a wavelength of 562nm with the Multiskan GO instrument and the Thermo Scientific SkanIt software (Thermo Scientific™, Waltham, Massachusetts, United States).

#### Western blot

Western blot analysis was performed for SEC fractions 2-9. An aliquot of 10µl (albumin) or 20 µl (ApoA1) was combined with a self-prepared 10X sample buffer and boiled for 10 min. Samples were loaded into 4–20% Mini-PROTEAN® TGX™ Precast Protein gels (#4561093, Bio-Rad Laboratories, Hercules, California, United States) and run at 200V for 35 min. The proteins were transferred to nitrocellulose membranes using Power Blotter Select Transfer Stacks (#PB3310, Invitrogen, Waltham, Massachusetts, USA) and semi dry transfer with Power Blotter Station (Invitrogen, Waltham, Massachusetts, USA). Membranes were blocked for 1h in TBS-Tween (0.1% Tween-20) 5% milk, and incubated overnight with primary antibodies against albumin (dilution 1:20,000, #ab194215, Abcam, Cambridge, UK) or ApoA1 (dilution 1:3000, #ab227455, Abcam, Cambridge, UK) in the blocking buffer. Membranes were washed three times and incubated with secondary antibodies: Donkey anti Sheep/Goat IgG (dilution 1:100,000, #STAR88P, Bio-Rad Laboratories, Hercules, California, United States), Goat Anti-Rabbit Immunoglobulins/HRP (dilution 1:100,000, #P0448, Dako Denmark A/S, Glostrup, Denmark) for 1h in RT, after which the washing steps were repeated. SuperSignal™ West Atto Ultimate Sensitivity Substrate (#A38554, Thermo Scientific™, Waltham, Massachusetts, United States) was applied on the membranes and the chemiluminescence was detected using FujiFilm LAS-3000 imager.

#### Nanoparticle tracking analysis

The samples were diluted in ultrapure water and the numbers of particles were measured using ZetaView® PMX120 (Particle Metrix, Meerbusch, Germany). Measurements were done at all 11 positions and the video quality was set to medium. The chamber temperature was set to 22°C, and the sensitivity of the camera to 85. Data were analysed using the ZetaView® analysis software version 8.05.12 with a minimum area of 10, a maximum area of 1000 and a minimum brightness of 30.

#### Nano-flow cytometry

A NanoAnalyzer U30 instrument (NanoFCM Co., Ltd, Nottingham, UK) with a 488nm laser and single-photon counting avalanche photodiode detectors (SPCM APDs) was used for determination of size and concentration of individual particles using side scattered light through a bandpass filter of 488/10 nm. The gravity-fed sheath fluid system consisted of HPLC grade water, focusing the sample core stream diameter to ∼1.4µm. Measurements were taken over 1-min durations at a sampling pressure of 1.0 kPa. Particle concentrations were determined against a standard of 250nm silica nanoparticles of known concentration. For particle sizing, a standard consisting of a 4-modal silica nanosphere cocktail (NanoFCM Inc., S16M-Exo) with diameters of 68, 91, 113 and 155nm was used. A standard curve was generated based on the side scattering intensity of the four different silica particle sub-populations using the NanoAnalyzer software. The laser was set to 10mW and 10% side scatter decay. Data processing and analysis was performed using the NanoFCM Professional Suite v2.0 software.

#### Transmission electron microscopy

The grids used were 200 square Mesh copper 3.05mm (G2200C, Agar Scientific, Stansted, UK). The film was made by hand in 2% biofoform in chloroform solution (SPI-Chem™ Pioloform® Resin, #2466, SPI Supplies, West Chester, PA, USA). Carbon coating was done with Bal-Tec CED030 carbon thread evaporator (Bal-Tec Union Ltd., Liechtenstein) and Emitech K100X Glow discharge unit (Emitech Ltd., UK) was used as the glow discharge system. Grids were then placed on a 12μl droplet of sample on parafilm for 2 minutes. Buffer salts were removed by transferring the grids twice to a fresh drop of distilled water and incubated for 5 s each. The excess fluid was removed with filter paper and the grids were transferred to one drop of uranyl acetate (1.5% in distilled water) and incubated for 1 minute. Excess fluid was removed with filter paper and the grids were air dried prior to imaging. The samples were imaged using a Jeol JEM-1400 transmission electron microscope with 80 000 V and magnifications of 6000X and 20000X.

#### Proteomics

EV samples were lysed, proteins precipitated and digested into peptides with trypsin using a previously published protocol for protein aggregation capture (Batth et al., 2019). In addition to fraction 3 (from 6ml, 3ml, 1ml, 0.5ml of CSF) and SEC fractions 4-5 (obtained from 6ml of CSF), also intact and depleted CSF samples were analysed with liquid chromatography with tandem mass spectrometry (LC-MS/MS). Intact CSF samples were prepared using the same protocol used with EVs. For depletion of the most abundant proteins from CSF ProteoSpin Abundant Serum Protein depletion kit (Norgen Biotek) was used according to manufacturer’s instructions, and proteins were in-solution digested with trypsin after depletion.

The resulting tryptic peptides were purified using home-made C18 Stage tips-columns followed by nanoLC-MS/MS analysis with 60-minute separation gradient using nanoElute coupled to timsTOF fleX (Bruker). For protein identification the data from LC-MS/MS was analysed by MaxQuant ver 2.0.1.0 against dog (Canis lupus) database downloaded from Uniprot. MaxQuant searches were done with and without ‘match between runs’ active.

The proteomics data are available via ProteomeXchange with identifier PXD036748.

#### Data analysis

In the proteomics data analysis, two different methods of identifying proteins were used: identification by LC-MS and identification by matching to a library created from all the analysed samples. For Venn diagrams, an online tool from Bioinformatics Institute Ghent was used (https://bioinformatics.psb.ugent.be/webtools/Venn/). Pathway analysis was done with FunRich version 3.1.4, with the level of significance set on p≤0.05.

Gene set enrichment analysis (GSEA) was performed with GSEA 4.1.0 program using ‘GSEApreranked’ option. A rank list was generated for each sample by giving the lowest rank number for protein with highest total intensity. After this an average of the ranks of three replicate samples was calculated and the final group rank list was generated based on that. Protein list for Vesiclepedia top 100 proteins was taken from its website on June 14^th^ 2022. Protein list for exomers was generated from differential expression data between small EVs and distinct nanoparticles from Zhang et al., 2019, proteins with positive logFC and adjusted p-value<0.05 were selected (Zhang et al., 2019). Out of the 103 proteins, 20 were identified from the proteomics data and were used as the protein list. Supermere protein list was generated by combining the top 20 most abundant proteins in supermeres derived from three cell lines presented in Supplementary Table 4 (Zhang et al., 2021). Out of the 34 proteins, 28 identified from the proteomics data were used as the protein list.

Proteomic and NTA data are presented as mean ± standard deviation unless otherwise stated.

## Results

### Literature review

At first, we performed a literature review to investigate the present state of EV analysis from CSF samples. A total of 122 articles were included in the review (**Supplementary file 1**). Of these, 104 (85%) studies used human CSF for studying the EVs whereas 16 (13%) studies used rodent CSF and 5 (4%) studies used CSF from other species such as pigs, sheep, and horses. In three (2%) of these articles both human and mouse CSF EVs were studied. The highest number of articles (n=34, 28% of total) focused on neurodegenerative diseases (NGD) such as Parkinson’s disease or Alzheimer’s disease (**Figure 1A**). The second highest group comprised articles related to immune-mediated demyelinating diseases (IMDDG), covering a total of 17 (14%) articles. Articles in this group included conditions such as multiple sclerosis or amyotrophic lateral sclerosis. Articles focusing on EV methodologies also formed a substantial set of 25 articles (21%). The ‘other’ group included other individual conditions, such as encephalitis, drug abuse, and exercise induced neuroprotection among others (n=19, 16%) (**Figure 1A**). Regarding the target of the analysis, most studies investigated EV protein content (n=52, 43%) either by analysing individual proteins (n=35, 29%) or with wider proteomic analysis (n=17, 14%). RNA content was studied in 48 articles (39%) and 18 articles (15%) focused on basic characterisation of EVs, including for example NTA or Western blot analysis. Only a few articles studied the biological function of CSF EVs (n=2, 2%) or lipidomics (n=2, 2%) (**Figure 1B**).

**Figure 1.**
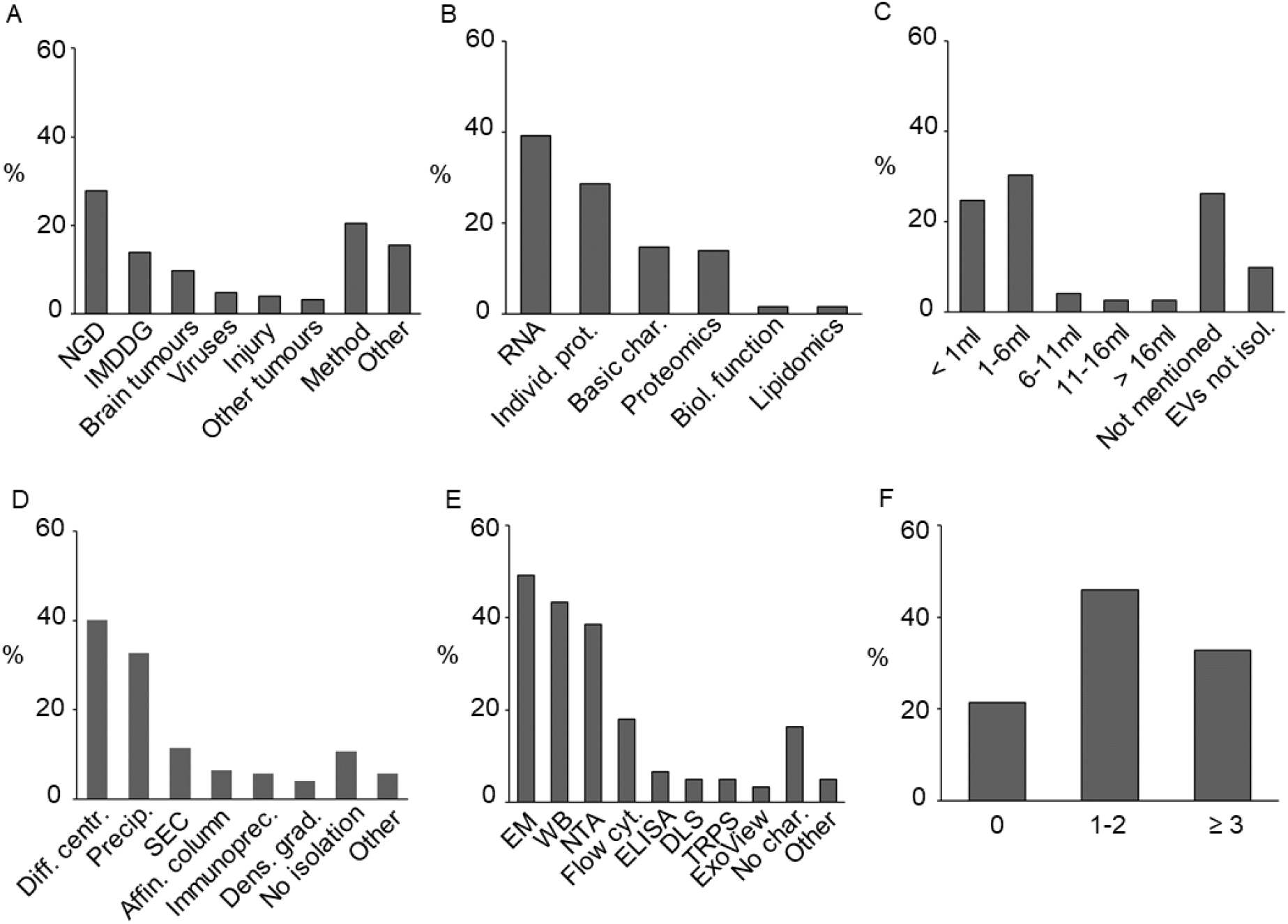
Data from the literature review of CSF EV articles published from 2015 to 2022. **A**. Conditions studied in the CSF EV articles included in this literature review. **B**. Targets of the CSF EV analyses. **C**. The starting volumes used prior to isolation of EVs. **D**. The isolation methods used. **E**. The EV characterisation methods used in the studies. **F**. Number of different characterisation methods used in the articles. Abbreviations: CSF = cerebrospinal fluid, EV = extracellular vesicle, NGD = neurodegenerative disease, IMDDG = immune-mediated demyelinating disease group, SEC = size- exclusion chromatography, WB = Western blot, NTA = nanoparticle tracking analysis, ELISA = Enzyme- Linked Immunosorbent Assay, DLS = dynamic light scattering, TRPS = tunable resistive pulse sensing.

Next, we reviewed the EV isolation and characterisation methods used in the articles and the volume of CSF from which EVs were derived. The highest number of studies (n=37, 30%) isolated EVs from 1-6ml of CSF, whereas 30 articles (25%) used starting volumes under 1ml. Unfortunately, a large number of studies (32 articles, 26%) did not report the starting volume (**Figure 1C**). Also, 12 articles (10%) did not use specific EV isolation methods despite studying EVs; in these, EVs were analysed directly from the CSF. Two isolation methods were used more often than the others: differential centrifugation-based methods in 49 articles (40%) and precipitation-based methods in 40 articles (33%) (**Figure 1D**). SEC-based methods were used in 14 articles (12%). Regarding the basic EV characterisation, the most commonly used methods were electron microscopy (EM) based techniques, Western blot, and NTA, which were used in 60 (49%), 53 (43%) and 47 (38%) articles, respectively (**Figure 1E)**. Most studies (56 articles, 46%) used one or two different methods to characterise the isolated EVs. In 40 articles (39%) three or more characterisation methods were used, however in 26 studies (21%) specific EV characterisation was not performed (**Figure 1F**).

A more detailed technical comparison was performed for the 17 articles which included proteomic analysis from CSF EVs (**Table 1**). A variety of EV isolation methods were used in these studies including differential centrifugation (n=4), precipitation-based methods (n=6) and SEC-based purification (n=5). The starting volume of CSF varied between 50μl and 8ml, but this did not clearly correlate with the number of proteins identified. Most of the articles used pooled CSF for analysis and only five articles analysed more than three biological replicates per group. Mass-spectrometry was the most commonly used technique, although commercial arrays (SOMAscan® array, PEA assay) were used in some articles.

**Table 1.**
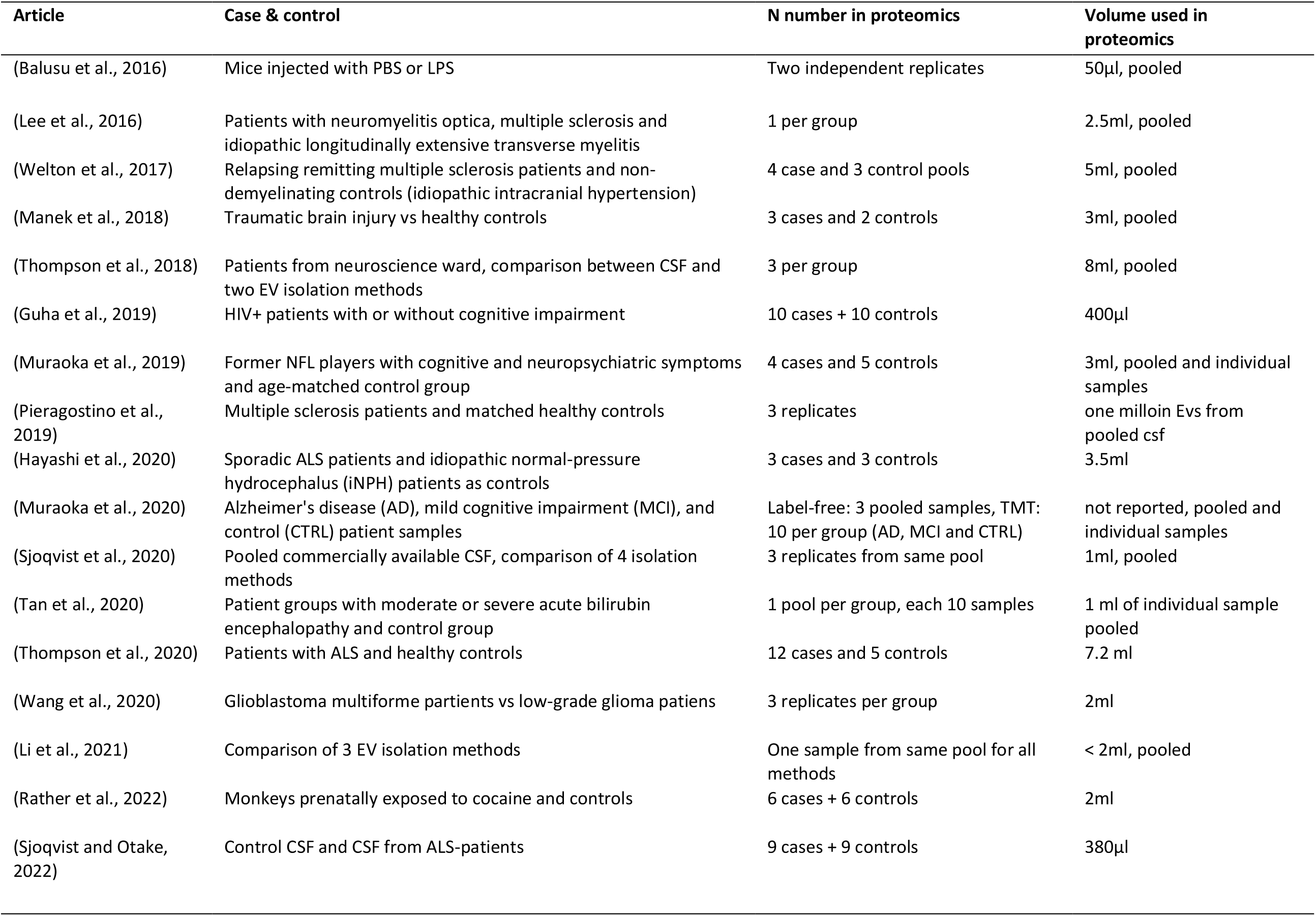
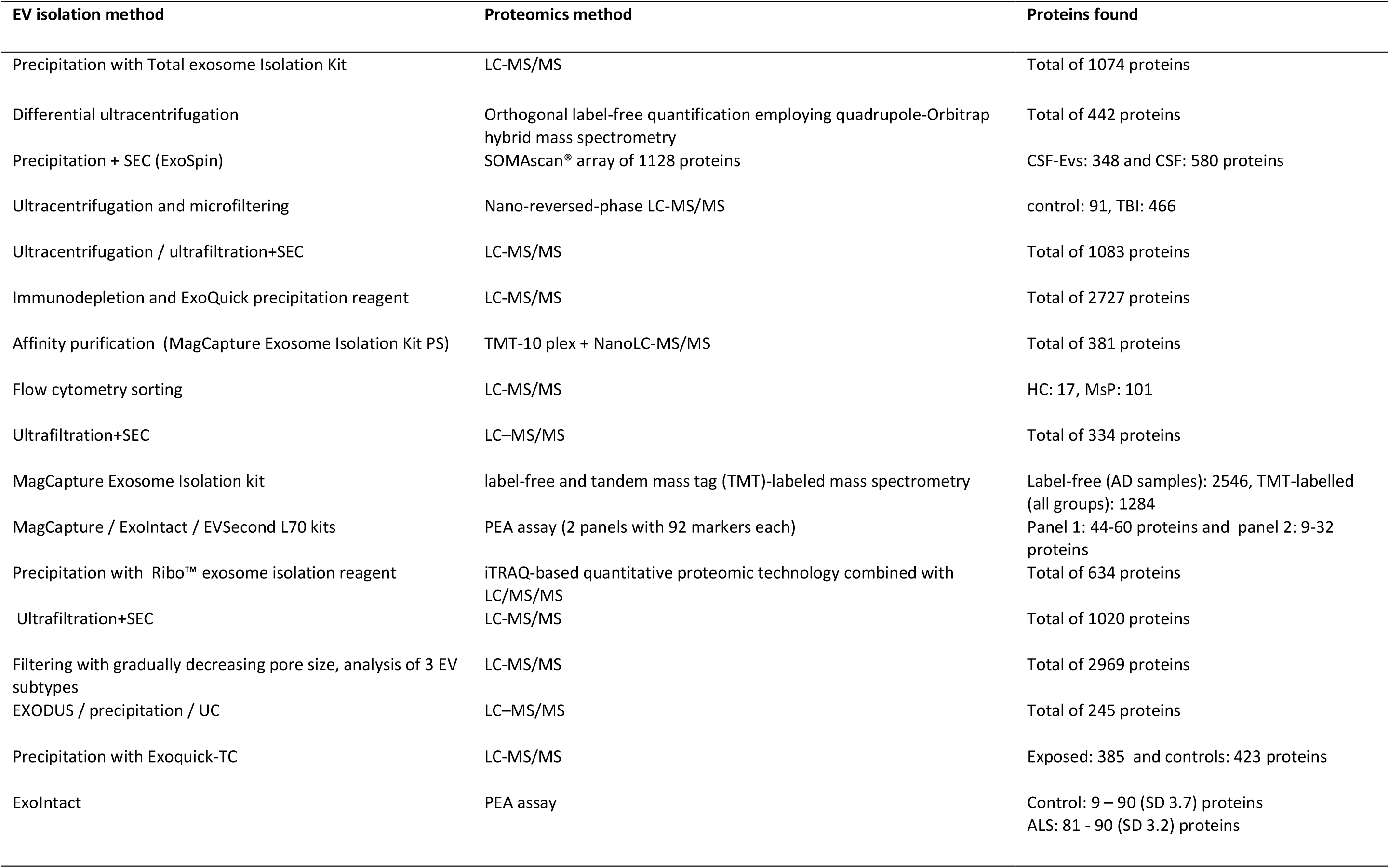
Review of articles including proteomics analysis from CSF-EVs.

### Sample characteristics

Samples from 21 dogs were included in this study. All but one dog had a CSF nucleated cell count of <5 cells/µl, and one sample had 10 cells/µl. All the CSF samples had a protein concentration within the reference interval (<300mg/l). Five separate CSF pools were used in the study, two for the method set-up and three for the proteomics analysis.

### EV isolation and characterisation from canine CSF

The EV characterisation was done with EVs derived from two separate 7ml pools of CSF. The highest number of particles (determined by NTA) was detected in fraction 3, as expected (**Figure 3A**). Protein concentration was highest in fractions 6-8 which was distinct from the particle peak (**Figure 2A**). The amount of protein in fractions 3 and 4 was under the detection threshold of the BCA assay and Western blot analysis was not sensitive enough to show the presence of canonical EV proteins (data not shown). Western blot analysis was performed as a purity control for two common EV-contamination proteins in CSF: albumin and ApoA1, a marker of high-density lipoproteins (HDL). Both of these proteins were present mainly in the protein fractions 6-7, confirming relatively good separation from the EVs (**Figure 2B**).

**Figure 2.**
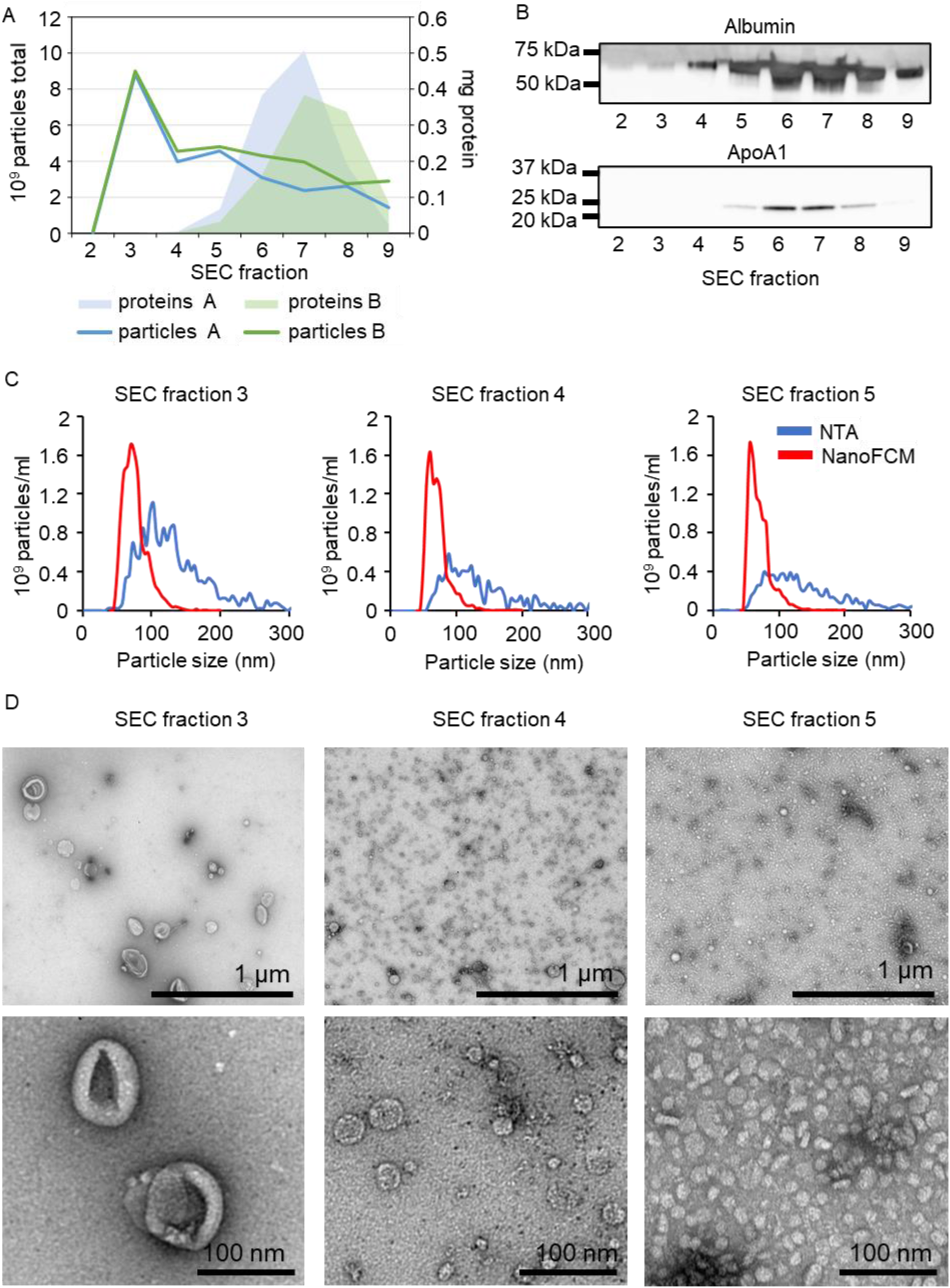
EV characterisation using UF-SEC. **A**. Total particle number measured with NTA and protein amount from SEC fractions 2-9 (each 500µl) from two separate CSF pools. **B**. Western blot analysis of albumin and ApoA1 from SEC fractions 2-9 from pool A. **C**. Particle size distribution measured with NTA and NanoFCM from SEC fractions 3-5 of pool B. **D**. TEM images from SEC fractions 3-5 from pool B. Abbreviations: EV = extracellular vesicle, UF-SEC = ultrafiltration combined with size-exclusion chromatography, CSF = cerebrospinal fluid, NTA = nanoparticle tracking analysis, NanoFCM = nano- flow cytometry, TEM = transmission electron microscopy.

**Figure 3.**
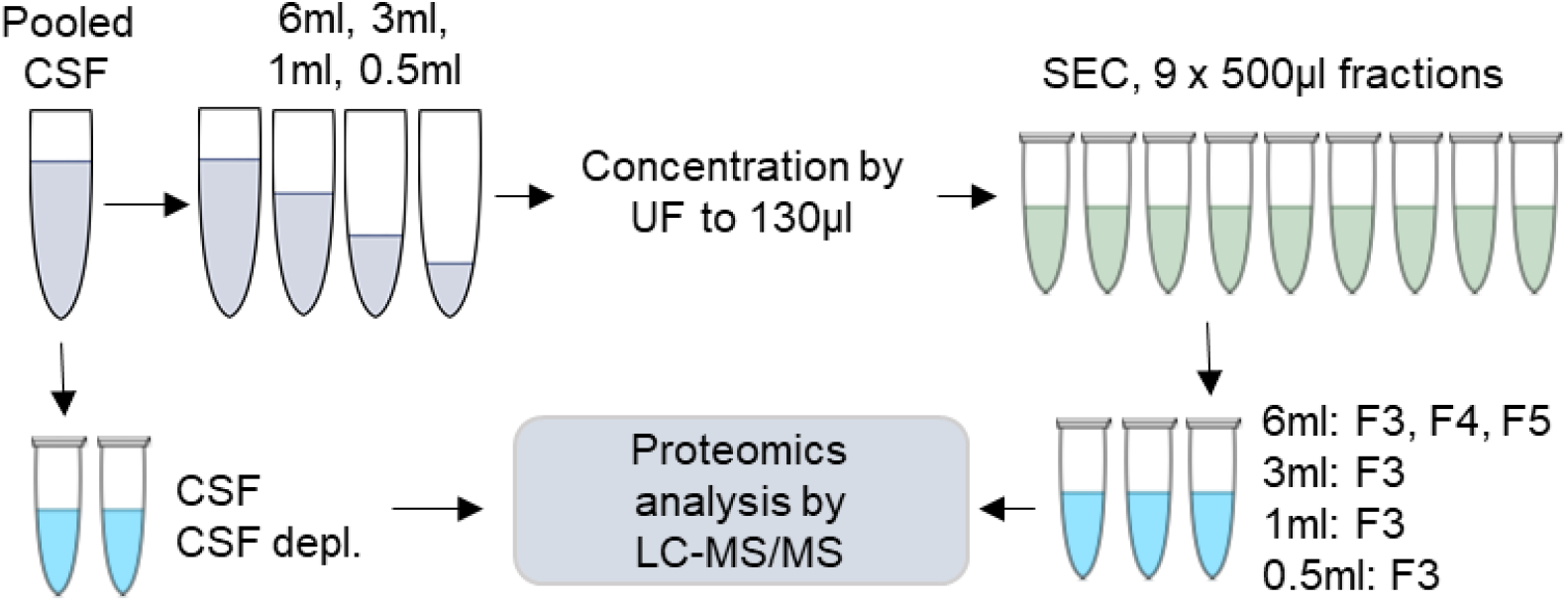
Workflow of the proteomic analysis. Pooled CSF was divided into a whole CSF sample with and without depletion, and into four different starting volumes (6ml, 3ml, 1ml, and 0.5ml). An UF-SEC protocol was used to fractionate the samples of the different volumes. At first, the CSF samples and fractions 3-5 from the 6ml starting volume were analysed and compared to evaluate the presence of EVs. In second comparison, the effect of CSF starting volume for EV protein contents was studied by comparing the fraction 3 of the four different starting volumes. Abbreviations: UF-SEC = ultrafiltration combined with size-exclusion chromatography, CSF: cerebrospinal fluid, EV = extracellular vesicle, CSF depl. = CSF with depletion of highest abundant plasma proteins, LC-MS/MS = liquid chromatography with tandem mass spectrometry.

The size distribution of the particles was measured with NTA and by nano-flow cytometry (NanoFCM). In general, NanoFCM showed a smaller particle diameter with a narrower particle size distribution compared to NTA. The particles in fraction 3 had a modal size of around 100nm when measured with NTA and around 70nm with NanoFCM. The measured particle size slightly decreased in fractions 4 and 5 with both analysis methods (**Figure 2C**).

Fractions 3-5 were further characterised with TEM and all of them had distinct particle populations. Fraction 3 included cup-shaped particles, a morphology commonly described for EVs (Lauer et al., 2016) (**Figure 2D**). The particle size was similar to NTA and NanoFCM data. Fraction 4 contained a main population of particles with a diameter of <50nm and fraction 5 even smaller particles, with a morphology resembling that of HDL (Lauer et al., 2016) (**Figure 2D**). Interestingly, most of the particles detected in fractions 4 and 5 were under the detection limit of NTA (70nm) and NanoFCM (45nm with scatter detection). In accordance with BCA results, fraction 5 included a darker cloud-like background considered most likely to reflect proteins.

### Proteomics comparison of SEC fractions and CSF

To further explore differences between the alleged EV fraction and the later SEC fractions, we compared the proteome of SEC fractions 3-5. The workflow for the proteomic analysis is illustrated in **Figure 3**. For identification, the LC-MS/MS data was analysed with MaxQuant search engine with and without ‘match between run (MBR)’ active. MBR increases the number of identifications for low-abundant proteins in the samples (Prianichnikov et al., 2020). We identified the highest number of proteins from fraction 3 (1493±107 and 999±173 proteins identified with and without MBR, respectively) and lowest number from whole CSF samples (1121±219 and 624±149 proteins identified with and without MBR, respectively) (**Figure 4A, Supplementary table1**).

**Figure 4.**
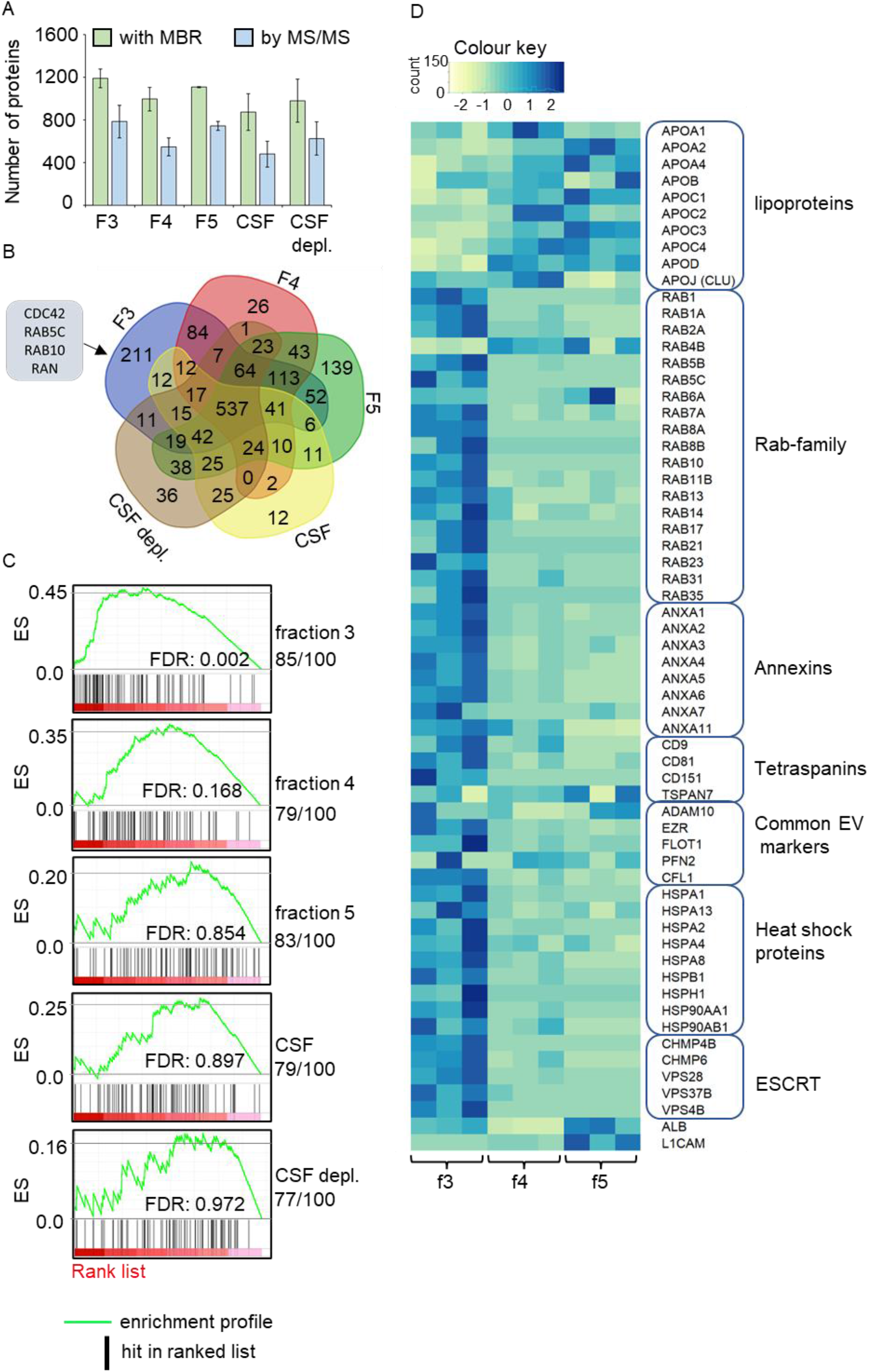
Proteomics comparison of SEC fractions and CSF. **A**. The number of identified proteins in the SEC fractions 3-5 isolated from 6ml of CSF, plain CSF and CSF depl. Protein number was calculated with and without the MBR setting. The values are given in mean and SD of three replicates. **B**. A Venn diagram from core protein list (found from each 3 rounds) of each fraction and CSF samples. Four proteins unique for fraction 3 aligning with the Vesiclepedia top 100 are listed. **C**. GSEA performed with the Vesiclepedia top 100 protein list. The number of proteins found from each rank list is given under the sample name. The ranked protein list was generated from the proteomics intensity data and is expressed in red in the plot, darker colour representing higher intensity. Black lines represent a protein included in the protein list and enrichment score represents the level of enrichment. **D**. Heatmap without clustering performed from iBAQ data for selected EV-related proteins. Abbreviations: CSF = cerebrospinal fluid, CSF depl. = CSF with depletion of highest abundant plasma proteins, MBR = matches between runs, SD = standard deviation, GSEA = Gene set enrichment analysis, iBAQ = Based Absolute Quantification, ES = enrichment profile, FDR = false recovery rate.

For the further comparisons, protein identification results using MBR were used. For each group (fractions 3-5, CSF, CSF depl.), a list of core proteins found in all three replicates were generated (**Supplementary figure 1**). When these core lists were compared with each other, 537 proteins were shared between all groups, and the highest number of unique proteins (n=211) was found from the EV-containing fraction 3, including four proteins (CDC42, RAB5C, RAB10, and RAN) from the Vesiclepedia top 100 list (**Figure 4B**). In CSF and CSF depl. combined, 73 proteins that were not present in the SEC fractions were identified. Also, 183 proteins that were not present in fraction 3 were identified in these two groups combined (**Figure 4B**).

Heatmap clustering performed for all identified proteins clustered samples from individual SEC fractions together and showed fraction-specific protein clusters **(Supplementary figure 2A)**. Especially fraction 3 included several clusters that were not abundant in other groups. In addition, the SEC fractions were compared with cellular component analysis using FunRich. Components considered significantly enriched in fraction 3 were exosomes, lysosome, and cytoplasm. Significantly enriched components for fraction 5 were endoplasmic reticulum and lysosome. No components were considered significant for fraction 4 **(Supplementary figure 2B)**.

Next, GSEA was used to probe the presence of EV and non-vesicular nanoparticle-related proteins in each fraction. In general, ranked lists for GSEA from all the samples identified ApoA1, ApoE and ApoJ (Clusterin) among the 6 highest ranked proteins, in other words they had high intensities. In addition, albumin was among the top 4 proteins in other groups apart from fraction 4, where it was ranked as the 11^th^ most abundant protein. Using Vesiclepedia top 100 proteins as a protein list, the greatest positive enrichment for EV associated proteins was detected within fraction 3 with a nominal p-value of <0.001 (**Figure 4C**). Fraction 4 was also enriched for EV associated proteins with a nominal p-value of 0.017, but with higher false discovery rate, indicating lower enrichment compared to fraction 3. Even though a relatively similar number of Vesiclepedia top 100 proteins were present in fraction 5 and whole CSF samples, they were not significantly enriched, and EV-related proteins had low relative intensity compared to other proteins in the samples (**Figure 4C**). Separate protein lists were generated for exomers and supermeres, recently reported non-vesicular nanoparticles with the size range present in TEM images from fractions 4 and 5 (Crescitelli et al., 2020; Zhang et al., 2021, 2019). With these lists, only enrichment detected was in fraction 3 using the supermere list. No enrichment was detected for the exomer list (**Supplementary figure 2C**).

As a final analysis, we compared the mean normalised Intensity Based Absolute Quantification (iBAQ) values from selected individual proteins from SEC fractions 3-5 **(Figure 4D)**. First group was apolipoproteins involved in the structures of HDL, low-density lipoproteins (LDL), chylomicrons, and ApoJ (Clusterin) which is connected to several neurodegenerative diseases (Yuste-Checa et al., 2022; Feingold, 2000). All apolipoproteins had highest intensities in fractions 4 and 5, indicating a presence of lipoprotein particles. To further explore EV-related proteins, we used a general EV marker listing prepared by Crescitelli et al., 2021 (Table 1 in their article) (Crescitelli et al., 2021). Apart from individual proteins (RAB4B, TSPAN7), the highest intensity was consistently detected in fraction 3. Lastly, albumin and L1-CAM were also added to the list as they are considered relevant regarding CSF EVs (Gomes and Witwer, 2022; Norman et al., 2021; Théry et al., 2018), both of which had highest intensity in fraction 5, likely reflecting the presence of non-vesicular proteins.

### Comparison of CSF starting volume

In addition to fraction comparison from 6ml of pooled CSF, UF-SEC isolation was performed from 3ml, 1ml and 0.5ml of each CSF pool. For those samples, proteomic analysis of fraction 3 was performed to compare the effect of starting volume on EV yield and number of proteins identified in the EV enriched fraction.

As expected, the highest total number of particles was found from the 6ml starting volume (10.6×10^9^±3.4×10^9^ particles) and the lowest in the 0.5ml starting volume (4.7×10^8^±1.7×10^8^ particles) **(Figure 5A)**. When particle number was normalised per 1ml of CSF used, the highest particle number was again found from the 6ml samples indicating a higher recovery of EVs from larger volumes compared to lower volumes. With the other starting volumes, the differences in EV recovery varied between rounds indicating no clear differences based on starting volume used **(Figure 5B)**.

**Figure 5.**
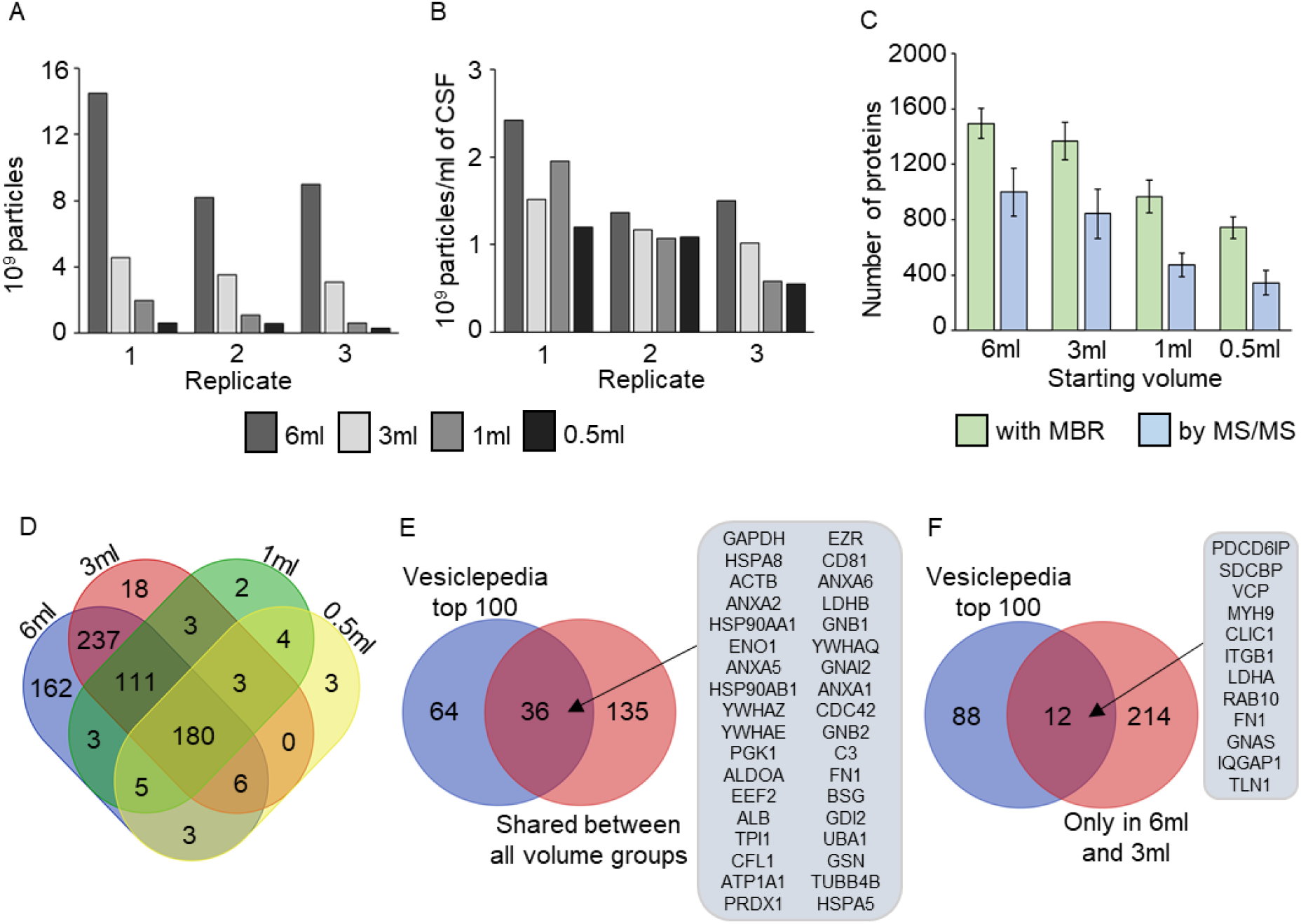
Comparison of CSF starting volume. **A**. The total number of particles in the EV fraction of each starting volume measured with NTA. **B**. The particle count per 1ml of CSF in the EV fraction of each starting volume. **C**. Total number of proteins identified from each starting volume’s EV fraction. The result is given as a mean and SD between all three replicates. **D**. A Venn diagram of the shared proteins between the different starting volumes’ EV fractions. Proteins identified in every replicate for each starting volume were included in the analysis. **E**. Alignment with the Vesiclepedia top 100 for the proteins shared between the EV fractions of all the starting volumes. Only proteins with corresponding gene names available were used for the analysis. The proteins aligning with Vesiclepedia are listed in the figure. **F**. Alignment with the Vesiclepedia top 100 for the proteins shared between the EV fractions of 6ml and 3ml starting volumes that were not identified in the 1ml or 0.5ml starting volumes. Only proteins with corresponding gene names available were used for the analysis. The proteins aligning with the Vesiclepedia top 100 are listed in the figure. Abbreviations: CSF = cerebrospinal fluid, EV = extracellular vesicle, NTA = nanoparticle tracking analysis, SD = standard deviation, MBR = matches between runs.

For the total number of proteins, the proteomics data was again analysed separately using both the MBR and ‘by MS-MS’ methods. With all samples, MBR analysis identified a higher number of proteins, with the highest number of proteins identified from the 6ml starting volume (1493±107 or 999±173 proteins) and the lowest number of proteins identified in the 0.5ml starting volume (743±77 or 345±88 proteins) (**Figure 5C**).

Further comparison of the starting volumes was done without MBR, as we were interested in finding out if the proteomic analysis can in the future be conducted on individual dogs without a large sample series. First, core lists of proteins derived from EVs isolated from different volumes of CSF were generated by listing all the proteins present in each proteomics replicate (**Supplementary figure 3A**). A Venn diagram of these core lists shows a total of 180 proteins present in all the volume groups (**Figure 5D**). For 171 of these, a gene name could be found (**Supplementary table 2). Of these 171 proteins, 36 were included in the Vesiclepedia top 100 list (Figure 5E**). In addition to the proteins shared between all the volume groups, the 6ml and 3ml volumes shared 237 proteins that were not present in the smaller volumes (**Figure 5D, Supplementary figure 3B**). Gene names were found for 226 of these, and 12 aligned with the Vesiclepedia top 100 (**Figure 5F**). A list of unique gene names of proteins identified in all three proteomics replicates in the 6ml starting volume is presented in **Supplementary table 3**, forming a list of stable EV-related proteins in canine CSF.

## Discussion

Here, our aim was to validate the UF-SEC method for isolation of EVs from canine CSF and to determine the smallest possible starting volume for reliable EV proteomic analysis. We mapped out the state of current CSF EV research and continued by comparing different SEC fractions and starting volumes of CSF.

The literature review revealed a high variability in the methods used and rigour in articles studying CSF EVs and that the reproducible information of EVs in CSF is still sparse. A large variety of different conditions has been studied, proving the potential for studying brain-related diseases. SEC is an increasingly popular technique for isolating the CSF EVs, however differential centrifugation or precipitation -based EV purification was used in over 70% of the articles, even though they are not optimal regarding the purity of vesicles (Clos-Sansalvador et al., 2022; Li et al., 2021). A variety of characterisation methods were used throughout the articles although electron microscopy, Western blot, and NTA were the most common. The majority of the articles characterised the EV preparations only with a maximum of two methods, and hence not fulfilling the MISEV guidelines (Théry et al., 2018). Furthermore, 11% of articles did not perform EV isolation and 21% did not characterise the EVs at all. This could be because collecting CSF is an invasive procedure and the volumes obtained are small, limiting the material available for basic characterisation. Thus, there is still a need for basic characterisation of CSF EVs and for the MISEV 2018 guidelines (Théry et al., 2018) to be followed more closely.

In our study, NTA measurement showed a clear particle peak at SEC fraction 3, which is consistent with findings when EVs are isolated from other body fluids using the same method (Karttunen et al., 2021). In fraction 3, particles detected by NTA had a modal size of around 100nm, whereas particles measured by NanoFCM had a modal size of around 70nm. Differences in the size distribution profile between these methods have also previously been reported (Arab et al., 2021).

Interestingly, TEM imaging also revealed distinct particle populations in fractions 4 and 5, with fraction 5 morphology resembling that of HDL (Lauer et al., 2016). Lipoprotein particles including HDL, LDL, VLDL and chylomicrons are known contaminants in EV preparations in plasma and they may co-isolate with EVs using SEC-based methods (Karimi et al., 2018; Simonsen, 2017). In addition, new types of smaller non-membranous extracellular particles have been discovered in recent years, including exomers and supermeres (Crescitelli et al., 2020; Zhang et al., 2021, 2019), and it could be hypothesised that these represent the particles seen in fractions 4 and 5. However, our data would not support this hypothesis since all proteins associated with small and large sized EVs (heat shock proteins were enriched in large EVs and tetraspanins, ADAMs, ESCRT proteins, SNAREs and Rab proteins were enriched in small EVs (Lischnig et al., 2022)) were found in fraction 3, and enrichment for supermere proteins was only identified in fraction 3. Also, the exomere listing was not enriched in fractions 3-5. Fractions 4 and 5 also had the highest relative intensity of apolipoproteins, supporting the premise that the particles identified in fractions 4 and 5 with TEM were mainly lipoproteins.

Despite separate protein profiles, the most abundant proteins were similar in all the SEC fractions and in whole CSF samples, and the number of shared proteins was high. Fraction 4 was enriched with Vesiclepedia top 100 proteins, but with lower abundance compared to fraction 3, whereas fraction 5 contained more non-vesicular proteins than the earlier SEC fractions. These data indicate that i) fraction 3 includes the majority of EVs, but also small quantities of lipoprotein particles, and ii) a small amount of EVs is present also in the other fractions. Although SEC did not fully fractionate EVs from other components of the CSF, fraction 3 did include the highest number of individual proteins compared to whole CSF, likely representing a reduction in the abundance of contaminant proteins (for example albumin) that might mask the presence of other proteins of lower abundance. Together with the Western blot analysis for ApoA1 and albumin, our proteomics data shows that our UF-SEC protocol was successful in reducing the quantity of non-vesicular proteins and lipoprotein contaminants from the EV fraction, which is in line with previous articles showing successful SEC-based EV isolation from CSF (Hayashi et al., 2020; Krušić Alić et al., 2022; Thompson et al., 2018).

L1-CAM has traditionally been considered as a marker for neuronal EVs (Gomes and Witwer, 2022), although it was recently suggested that L1-CAM or its isoforms are not carried by EVs in either human plasma or CSF (Norman et al., 2021). In our study, L1-CAM was identified from the unpurified CSF and from the SEC fractions 4 and 5, and was not consistently identified within fraction 3. These data indicate that L1-CAM is not associated with EVs in canine CSF, which supports the data from the aforementioned study (Norman et al., 2021).

The previous proteomics studies from CSF EVs have used variable starting volumes and methods for EV isolation, and most used pooled CSF. Pooled samples are useful in methodological comparisons, but in the clinical setting analysis of individual patient samples is usually required, although the limited CSF volume that can be obtained from patients ante-mortem can be a limiting factor. As expected, both absolute EV yield and the number of proteins identified correlated with the starting volume used. Promisingly, sufficient amount of EVs could be isolated from 0.5ml of CSF to permit proteomic analysis, with 293-447 proteins identified from such samples without the MBR setting active, thus providing proof of concept that proteomic analysis of EVs could be performed on individual canine patient samples. In addition, when the MaxQuant analysis was performed using the MBR setting, we were able to identify 696-832 proteins from the 0.5ml samples. This could be a useful option for further clinical proteomics studies on CSF EVs if it is possible to include one sample with larger starting volume to boost the overall number of identifications.

Despite the challenges, CSF EVs hold high potential for biomarker studies of brain related diseases. There is an ever-growing interest in EV research, but CSF EVs are yet to be studied on a larger scale. Our data suggest that the UF-SEC method effectively enriches EVs and reduces the amount of contaminant proteins present in EV enriched fractions. The NTA and proteomic analysis showed that sufficient EVs can be isolated from 0.5ml of CSF to permit proteomic analysis, although a volume of ≥3ml of CSF is likely to allow more complete proteomic analysis.

### Geolocation information

The study was conducted in the University of Helsinki, Finland. We collaborated with the University of Oslo and Oslo University Hospital in Norway, Helsinki University Hospital in Finland, with NanoFCM co., Ltd in the United Kingdom, and with University of Cambridge in the United Kingdom.

## Acknowledgements

This work was supported by the Academy of Finland under Grant number 332789; and Finnish Foundation of Veterinary Research with a scholarship. Mass spectrometry-based proteomic analyses were performed by the Proteomics Core Facility, Department of Immunology, University of Oslo/Oslo University Hospital, which is supported by the Core Facilities program of the South-Eastern Norway Regional Health Authority. This core facility is also a member of the National Network of Advanced Proteomics Infrastructure (NAPI), which is funded by the Research Council of Norway INFRASTRUKTUR-program (project number: 295910). Nanoparticle tracking analysis was performed at the University of Helsinki EV core, and TEM imaging at the Helsinki Electron Microscopy Core Unit. NanoFCM analysis was done at NanoFCM co., Ltd. We also thank Ninna Koho for helping us with the laboratory work.

## Author contribution

Conseptualisation: Petra Kangas, Jenni Karttunen, Tarja S. Jokinen, Tuula A. Nyman, Tim Williams, Liisa Metsähonkala. Methodology: Petra Kangas, Jenni Karttunen, Tarja S. Jokinen, Tuula A. Nyman, Cameron Burns, and Robert Tempest. Data analysis and writing the original manuscript: Petra Kangas and Jenni Karttunen. Writing (review and editing): Petra Kangas, Jenni Karttunen, Tarja S. Jokinen, Tuula A. Nyman, Cameron Burns, Robert Tempest, Tim Williams, and Liisa Metsähonkala. Visualisation: Petra Kangas and Jenni Karttunen. Project administration and funding acquisition: Tarja S. Jokinen. All authors have read and agreed to the published version of the manuscript.

