## Supplementary file 1 for "Towards optimised extracellular vesicle proteomics from cerebrospinal fluid"

*Shared last author

**Supplementary file 1:** Articles analysed for the literature review.

Akers, Johnny C., Wei Hua, Hongying Li, Valya Ramakrishnan, Zixiao Yang, Kai Quan, Wei Zhu, et al. 'A Cerebrospinal Fluid MicroRNA Signature as Biomarker for Glioblastoma'. *Oncotarget* 8, no. 40 (15 September 2017): 68769–79. https://doi.org/10.18632/oncotarget.18332.

Akers, Johnny C., Valya Ramakrishnan, Ryan Kim, Shirley Phillips, Vivek Kaimal, Ying Mao, Wei Hua, et al. 'MiRNA Contents of Cerebrospinal Fluid Extracellular Vesicles in Glioblastoma Patients'. *Journal of Neuro-Oncology* 123, no. 2 (June 2015): 205–16. https://doi.org/10.1007/s11060-015-1784-3.

Akers, Johnny C., Valya Ramakrishnan, John P. Nolan, Erika Duggan, Chia-Chun Fu, Fred H. Hochberg, Clark C. Chen, and Bob S. Carter. 'Comparative Analysis of Technologies for Quantifying Extracellular Vesicles (EVs) in Clinical Cerebrospinal Fluids (CSF)'. Edited by Hang Hubert Yin. *PLOS ONE* 11, no. 2 (22 February 2016): e0149866. https://doi.org/10.1371/journal.pone.0149866.

Akers, Johnny C., Valya Ramakrishnan, Isaac Yang, Wei Hua, Ying Mao, Bob S. Carter, and Clark C. Chen. 'Optimizing Preservation of Extracellular Vesicular MiRNAs Derived from Clinical Cerebrospinal Fluid'. *Cancer Biomarkers* 17, no. 2 (3 August 2016): 125–32. https://doi.org/10.3233/CBM-160609.

Alsop, Eric, Bessie Meechoovet, Robert Kitchen, Thadryan Sweeney, Thomas G. Beach, Geidy E. Serrano, Elizabeth Hutchins, et al. 'A Novel Tissue Atlas and Online Tool for the Interrogation of Small RNA Expression in Human Tissues and Biofluids'. *Frontiers in Cell and Developmental Biology* 10 (4 March 2022): 804164. https://doi.org/10.3389/fcell.2022.804164.

Anderson, Monique R., Michelle L. Pleet, Yoshimi Enose-Akahata, James Erickson, Maria Chiara Monaco, Yao Akpamagbo, Ashley Velluci, et al. 'Viral Antigens Detectable in CSF Exosomes from Patients with Retrovirus Associated Neurologic Disease: Functional Role of Exosomes'. *Clinical and Translational Medicine* 7, no. 1 (December 2018). https://doi.org/10.1186/s40169-018-0204-7.

Balusu, Sriram, Elien Van Wonterghem, Riet De Rycke, Koen Raemdonck, Stephan Stremersch, Kris Gevaert, Marjana Brkic, et al. 'Identification of a Novel Mechanism of Blood–Brain Communication during Peripheral Inflammation via Choroid Plexus-derived Extracellular Vesicles'. *EMBO Molecular Medicine* 8, no. 10 (October 2016): 1162–83. https://doi.org/10.15252/emmm.201606271.

Castañeyra-Ruiz, Leandro, Ibrahim González-Marrero, Luis G. Hernández-Abad, Emilia M. Carmona-Calero, Marta R. Pardo, Rebeca Baz-Davila, Seunghyun Lee, Michael Muhonen, Ricardo Borges, and Agustín Castañeyra-Perdomo. 'AQP4 Labels a Subpopulation of White Matter-Dependent Glial Radial Cells Affected by Pediatric Hydrocephalus, and Its Expression Increased in Glial Microvesicles Released to the Cerebrospinal Fluid in Obstructive Hydrocephalus'. *Acta Neuropathologica Communications* 10, no. 1 (December 2022): 41. https://doi.org/10.1186/s40478-022-01345-4.

Cheng, Peng, Feifei Feng, Hui Yang, Suqin Jin, Chao Lai, Yun Wang, and Jianzhong Bi. 'Detection and Significance of Exosomal MRNA Expression Profiles in the Cerebrospinal Fluid of Patients with Meningeal Carcinomatosis'. *Journal of Molecular Neuroscience* 71, no. 4 (April 2021): 790–803. https://doi.org/10.1007/s12031-020-01701-w.

Colombo, Federico, Mattia Bastoni, Annamaria Nigro, Paola Podini, Annamaria Finardi, Giacomo Casella, Menon Ramesh, Cinthia Farina, Claudia Verderio, and Roberto Furlan. 'Cytokines Stimulate the Release of Microvesicles from Myeloid Cells Independently from the P2X7 Receptor/Acid Sphingomyelinase Pathway'. *Frontiers in Immunology* 9 (7 February 2018): 204. https://doi.org/10.3389/fimmu.2018.00204.

Costa, Júlia, Ana Pronto-Laborinho, Susana Pinto, Marta Gromicho, Sara Bonucci, Erin Tranfield, Catarina Correia, Bruno M. Alexandre, and Mamede de Carvalho. 'Investigating LGALS3BP/90 K Glycoprotein in the Cerebrospinal Fluid of Patients with Neurological Diseases'. *Scientific Reports* 10, no. 1 (December 2020): 5649. https://doi.org/10.1038/s41598-020-62592-w.

Cressatti, Marisa, Julia M. Galindez, Lamin Juwara, Natalie Orlovetskie, Ana M. Velly, Shaun Eintracht, Adrienne Liberman, Mervyn Gornitsky, and Hyman M. Schipper. 'Characterization and Heme Oxygenase-1 Content of Extracellular Vesicles in Human Biofluids'. *Journal of Neurochemistry* 157, no. 6 (June 2021): 2195–2209. https://doi.org/10.1111/jnc.15167.

Crotti, Andrea, Hameetha Rajamohamend Sait, Kathleen M. McAvoy, Karol Estrada, Ayla Ergun, Suzanne Szak, Galina Marsh, et al. 'BIN1 Favors the Spreading of Tau via Extracellular Vesicles'. *Scientific Reports* 9, no. 1 (December 2019): 9477. https://doi.org/10.1038/s41598-019-45676-0.

Cruz, Allecineia Bispo da, Marta Marques Maia, Ingrid de Siqueira Pereira, Noemi Nosomi Taniwaki, Gislene Mitsue Namiyama, João Paulo Marochi Telles, Jose Ernesto Vidal, et al. 'Human Extracellular Vesicles and Correlation with Two Clinical Forms of Toxoplasmosis'. Edited by Gordon Langsley. *PLOS ONE* 15, no. 3 (3 March 2020): e0229602. https://doi.org/10.1371/journal.pone.0229602.

Daaboul, George G., Paola Gagni, Luisa Benussi, Paolo Bettotti, Miriam Ciani, Marina Cretich, David S. Freedman, et al. 'Digital Detection of Exosomes by Interferometric Imaging'. *Scientific Reports* 6, no. 1 (December 2016): 37246. https://doi.org/10.1038/srep37246.

Dalla Costa, Gloria, Tommaso Croese, Marco Pisa, Annamaria Finardi, Lorena Fabbella, Vittorio Martinelli, Letizia Leocani, Massimo Filippi, Giancarlo Comi, and Roberto Furlan. 'CSF Extracellular Vesicles and Risk of Disease Activity after a First Demyelinating Event'. *Multiple Sclerosis Journal* 27, no. 10 (September 2021): 1606–10. https://doi.org/10.1177/1352458520987542.

Derkow, Katja, Rosa Rössling, Carola Schipke, Christina Krüger, Jakob Bauer, Michael Fähling, Andrea Stroux, et al. 'Distinct Expression of the Neurotoxic MicroRNA Family Let-7 in the Cerebrospinal Fluid of Patients with Alzheimer's Disease'. Edited by Jing A. Zhang. *PLOS ONE* 13, no. 7 (16 July 2018): e0200602. https://doi.org/10.1371/journal.pone.0200602.

Ding, Xuebing, Mingming Ma, Junfang Teng, Robert K.F. Teng, Shuang Zhou, Jingzheng Yin, Ekokobe Fonkem, Jason H. Huang, Erxi Wu, and Xuejing Wang. 'Exposure to ALS-FTD-CSF Generates TDP-43 Aggregates in Glioblastoma Cells through Exosomes and TNTs-like Structure'. *Oncotarget* 6, no. 27 (15 September 2015): 24178–91. https://doi.org/10.18632/oncotarget.4680.

Dozio, Vito, Veerle Lejon, Dieudonné Mumba Ngoyi, Philippe Büscher, Jean-Charles Sanchez, and Natalia Tiberti. 'Cerebrospinal Fluid-Derived Microvesicles From Sleeping Sickness Patients Alter Protein Expression in Human Astrocytes'. *Frontiers in Cellular and Infection Microbiology* 9 (20 November 2019): 391. https://doi.org/10.3389/fcimb.2019.00391.

Egyed, Bálint, Nóra Kutszegi, Judit C. Sági, András Gézsi, Andrea Rzepiel, Tamás Visnovitz, Péter Lőrincz, et al. 'MicroRNA-181a as Novel Liquid Biopsy Marker of Central Nervous System Involvement in Pediatric Acute Lymphoblastic Leukemia'. *Journal of Translational Medicine* 18, no. 1 (December 2020): 250. https://doi.org/10.1186/s12967-020-02415-8.

Eitan, Erez, Emmette R Hutchison, Krisztina Marosi, James Comotto, Maja Mustapic, Saket M Nigam, Caitlin Suire, et al. 'Extracellular Vesicle-Associated Aβ Mediates Trans-Neuronal Bioenergetic and Ca2+-Handling Deficits in Alzheimer's Disease Models'. *Npj Aging and Mechanisms of Disease* 2, no. 1 (December 2016): 16019. https://doi.org/10.1038/npjamd.2016.19.

Elkouris, Maximilianos, Georgia Kouroupi, Alexios Vourvoukelis, Nikolaos Papagiannakis, Valeria Kaltezioti, Rebecca Matsas, Leonidas Stefanis, Maria Xilouri, and Panagiotis K. Politis. 'Long Non-Coding RNAs Associated With Neurodegeneration-Linked Genes Are Reduced in Parkinson's Disease Patients'. *Frontiers in Cellular Neuroscience* 13 (22 February 2019): 58. https://doi.org/10.3389/fncel.2019.00058.

Emelyanov, Anton, Tatiana Shtam, Roman Kamyshinsky, Luiza Garaeva, Nikolai Verlov, Irina Miliukhina, Anastasia Kudrevatykh, et al. 'Cryo-Electron Microscopy of Extracellular Vesicles from Cerebrospinal Fluid'. Edited by Giovanni Camussi. *PLOS ONE* 15, no. 1 (30 January 2020): e0227949. https://doi.org/10.1371/journal.pone.0227949.

Figueroa, Javier M, Johan Skog, Johnny Akers, Hongying Li, Ricardo Komotar, Randy Jensen, Florian Ringel, et al. 'Detection of Wild-Type EGFR Amplification and EGFRvIII Mutation in CSF-Derived Extracellular Vesicles of Glioblastoma Patients'. *Neuro-Oncology* 19, no. 11 (19 October 2017): 1494–1502. https://doi.org/10.1093/neuonc/nox085.

Galazka, Grazyna, Marcin P Mycko, Igor Selmaj, Cedric S Raine, and Krzysztof W Selmaj. 'Multiple Sclerosis: Serum-Derived Exosomes Express Myelin Proteins'. *Multiple Sclerosis Journal* 24, no. 4 (April 2018): 449–58. https://doi.org/10.1177/1352458517696597.

Gelibter, Stefano, Marco Pisa, Tommaso Croese, Annamaria Finardi, Alessandra Mandelli, Francesca Sangalli, Bruno Colombo, et al. 'Spinal Fluid Myeloid Microvesicles Predict Disease Course in Multiple Sclerosis'. *Annals of Neurology* 90, no. 2 (August 2021): 253–65. https://doi.org/10.1002/ana.26154.

Geraci, Fabiana, Paolo Ragonese, Maria Magdalena Barreca, Emanuele Aliotta, Maria Antonietta Mazzola, Sabrina Realmuto, Giulia Vazzoler, Giovanni Savettieri, Gabriella Sconzo, and Giuseppe Salemi. 'Differences in Intercellular Communication During Clinical Relapse and Gadolinium-Enhanced MRI in Patients With Relapsing Remitting Multiple Sclerosis: A Study of the Composition of Extracellular Vesicles in Cerebrospinal Fluid'. *Frontiers in Cellular Neuroscience* 12 (15 November 2018): 418. https://doi.org/10.3389/fncel.2018.00418.

Gomes de Andrade, Gisele, Laura Reck Cechinel, Karine Bertoldi, Fernando Galvão, Paulo Valdeci Worm, and Ionara Rodrigues Siqueira. 'The Aging Process Alters IL-1β and CD63 Levels

Differently in Extracellular Vesicles Obtained from the Plasma and Cerebrospinal Fluid'. *Neuroimmunomodulation* 25, no. 1 (2018): 18–22. https://doi.org/10.1159/000488943.

Goswami, Saptamita, Atoshi Banerjee, Bharti Kumari, Bhaswati Bandopadhyay, Nemai Bhattacharya, Nandita Basu, Sudhanshu Vrati, and Arup Banerjee. 'Differential Expression and Significance of Circulating MicroRNAs in Cerebrospinal Fluid of Acute Encephalitis Patients Infected with Japanese Encephalitis Virus'. *Molecular Neurobiology* 54, no. 2 (March 2017): 1541–51. https://doi.org/10.1007/s12035-016-9764-y.

Gu, Jiachen, Tao Jin, Zongshan Li, Huimin Chen, Hongbo Xia, Xiaomin Xu, and Yaxing Gui. 'Exosomes Expressing Neuronal Autoantigens Induced Immune Response in Antibody-Positive Autoimmune Encephalitis'. *Molecular Immunology* 131 (March 2021): 164–70. https://doi.org/10.1016/j.molimm.2020.12.034.

Guha, Debjani, David R. Lorenz, Vikas Misra, Sukrutha Chettimada, Susan Morgello, and Dana Gabuzda. 'Proteomic Analysis of Cerebrospinal Fluid Extracellular Vesicles Reveals Synaptic Injury, Inflammation, and Stress Response Markers in HIV Patients with Cognitive Impairment'. *Journal of Neuroinflammation* 16, no. 1 (December 2019): 254. https://doi.org/10.1186/s12974-019-1617-y.

Guha, Debjani, Shibani S. Mukerji, Sukrutha Chettimada, Vikas Misra, David R. Lorenz, Susan Morgello, and Dana Gabuzda. 'Cerebrospinal Fluid Extracellular Vesicles and Neurofilament Light Protein as Biomarkers of Central Nervous System Injury in HIV-Infected Patients on Antiretroviral Therapy'. *AIDS* 33, no. 4 (15 March 2019): 615–25. https://doi.org/10.1097/QAD.0000000000002121.

Gui, YaXing, Hai Liu, LiShan Zhang, Wen Lv, and XingYue Hu. 'Altered MicroRNA Profiles in Cerebrospinal Fluid Exosome in Parkinson Disease and Alzheimer Disease'. *Oncotarget* 6, no. 35 (10 November 2015): 37043–53. https://doi.org/10.18632/oncotarget.6158.

Guix, Francesc, Grant Corbett, Diana Cha, Maja Mustapic, Wen Liu, David Mengel, Zhicheng Chen, et al. 'Detection of Aggregation-Competent Tau in Neuron-Derived Extracellular Vesicles'. *International Journal of Molecular Sciences* 19, no. 3 (27 February 2018): 663. https://doi.org/10.3390/ijms19030663.

Hayashi, Noriko, Hiroshi Doi, Yoichi Kurata, Hiroyuki Kagawa, Yoshitoshi Atobe, Kengo Funakoshi, Mikiko Tada, et al. 'Proteomic Analysis of Exosome-Enriched Fractions Derived from Cerebrospinal Fluid of Amyotrophic Lateral Sclerosis Patients'. *Neuroscience Research* 160 (November 2020): 43–49. https://doi.org/10.1016/j.neures.2019.10.010.

He, Jinting, Ming Ren, Haiqi Li, Le Yang, Xiaofeng Wang, and Qiwei Yang. 'Exosomal Circular RNA as a Biomarker Platform for the Early Diagnosis of Immune-Mediated Demyelinating Disease'. *Frontiers in Genetics* 10 (27 September 2019): 860. https://doi.org/10.3389/fgene.2019.00860.

Henderson, Lisa J., Tory P. Johnson, Bryan R. Smith, Lauren Bowen Reoma, Ulisses A. Santamaria, Muzna Bachani, Catherine Demarino, et al. 'Presence of Tat and Transactivation Response Element in Spinal Fluid despite Antiretroviral Therapy'. *AIDS* 33, no. Supplement 2 (1 December 2019): S145–57. https://doi.org/10.1097/QAD.0000000000002268.

Hong, Zhen, Chen Tian, Tessandra Stewart, Patrick Aro, David Soltys, Matt Bercow, Lifu Sheng, et al. 'Development of a Sensitive Diagnostic Assay for Parkinson Disease Quantifying α-Synuclein–Containing Extracellular Vesicles'. *Neurology* 96, no. 18 (4 May 2021): e2332–45. https://doi.org/10.1212/WNL.0000000000011853.

Hou, Xiaocan, Xuan Gong, Longbo Zhang, Tianjiao Li, Hongyu Yuan, Yue Xie, Yun Peng, et al. 'Identification of a Potential Exosomal Biomarker in Spinocerebellar Ataxia Type 3/Machado–

Joseph Disease'. *Epigenomics* 11, no. 9 (July 2019): 1037–56. https://doi.org/10.2217/epi-2019-0081.
Huang, Mudan, Chongjun Xiao, Liying Zhang, Lili Li, Jing Luo, Lilin Chen, Xiquan Hu, and Haiqing Zheng. 'Bioinformatic Analysis of Exosomal MicroRNAs of Cerebrospinal Fluid in Ischemic Stroke Rats After Physical Exercise'. *Neurochemical Research* 46, no. 6 (June 2021): 1540–53. https://doi.org/10.1007/s11064-021-03294-1.
Jain, Gaurav, Anne Stuendl, Pooja Rao, Tea Berulava, Tonatiuh Pena Centeno, Lalit Kaurani, Susanne Burkhardt, et al. 'A Combined MiRNA–PiRNA Signature to Detect Alzheimer's Disease'. *Translational Psychiatry* 9, no. 1 (December 2019): 250. https://doi.org/10.1038/s41398-019-0579-2.
Kim, Seh Hyun, Sin-Weon Yun, Hye Ryoun Kim, and Soo Ahn Chae. 'Exosomal MicroRNA Expression Profiles of Cerebrospinal Fluid in Febrile Seizure Patients'. *Seizure* 81 (October 2020): 47–52. https://doi.org/10.1016/j.seizure.2020.07.015.
Kong, Fan-Long, Xiao-Ping Wang, Ya-Nan Li, and Hai-Xu Wang. 'The Role of Exosomes Derived from Cerebrospinal Fluid of Spinal Cord Injury in Neuron Proliferation *in Vitro*'. *Artificial Cells, Nanomedicine, and Biotechnology* 46, no. 1 (2 January 2018): 200–205. https://doi.org/10.1080/21691401.2017.1304408.
Krušić Alić, Vedrana, Mladenka Malenica, Maša Biberić, Siniša Zrna, Lara Valenčić, Aleksandar Šuput, Lada Kalagac Fabris, et al. 'Extracellular Vesicles from Human Cerebrospinal Fluid Are Effectively Separated by Sepharose CL-6B—Comparison of Four Gravity-Flow Size Exclusion Chromatography Methods'. *Biomedicines* 10, no. 4 (27 March 2022): 785. https://doi.org/10.3390/biomedicines10040785.
Kuharić, Janja, Kristina Grabušić, Vlatka Sotošek Tokmadžić, Sanja Štifter, Ksenija Tulić, Olga Shevchuk, Pero Lučin, and Alan Šustić. 'Severe Traumatic Brain Injury Induces Early Changes in the Physical Properties and Protein Composition of Intracranial Extracellular Vesicles'. *Journal of Neurotrauma* 36, no. 2 (15 January 2019): 190–200. https://doi.org/10.1089/neu.2017.5515.
Kurzawa-Akanbi, Marzena, Seshu Tammireddy, Ivo Fabrik, Lina Gliaudelytė, Mary K. Doherty, Rachel Heap, Irena Matečko-Burmann, et al. 'Altered Ceramide Metabolism Is a Feature in the Extracellular Vesicle-Mediated Spread of Alpha-Synuclein in Lewy Body Disorders'. *Acta Neuropathologica* 142, no. 6 (December 2021): 961–84. https://doi.org/10.1007/s00401-021-02367-3.
Lee, Jingyun, Kimberly Q. McKinney, Antonis J. Pavlopoulos, May H. Han, Su-Hyun Kim, Ho Jin Kim, and Sunil Hwang. 'Exosomal Proteome Analysis of Cerebrospinal Fluid Detects Biosignatures of Neuromyelitis Optica and Multiple Sclerosis'. *Clinica Chimica Acta* 462 (November 2016): 118–26. https://doi.org/10.1016/j.cca.2016.09.001.
Lee, Kyue-Yim, Ji Hye Im, Weiwei Lin, Ho-Shin Gwak, Jong Heon Kim, Byong Chul Yoo, Tae Hoon Kim, et al. 'Nanoparticles in 472 Human Cerebrospinal Fluid: Changes in Extracellular Vesicle Concentration and MiR-21 Expression as a Biomarker for Leptomeningeal Metastasis'. *Cancers* 12, no. 10 (24 September 2020): 2745. https://doi.org/10.3390/cancers12102745.
Rivero Vaccari, Juan Pablo de, Frank Brand, Stephanie Adamczak, Stephanie W. Lee, Jon Perez-Barcena, Michael Y. Wang, M. Ross Bullock, W. Dalton Dietrich, and Robert W. Keane. 'Exosome-Mediated Inflammasome Signaling after Central Nervous System Injury'. *Journal of Neurochemistry* 136 (January 2016): 39–48. https://doi.org/10.1111/jnc.13036.
Lee, Kyue-Yim, Yoona Seo, Ji Hye Im, Jiho Rhim, Woosun Baek, Sewon Kim, Ji-Woong Kwon, et al. 'Molecular Signature of Extracellular Vesicular Small Non-Coding RNAs Derived from

Cerebrospinal Fluid of Leptomeningeal Metastasis Patients: Functional Implication of MiR-21 and Other Small RNAs in Cancer Malignancy'. *Cancers* 13, no. 2 (8 January 2021): 209. https://doi.org/10.3390/cancers13020209.

Lepko, Tjasa, Melanie Pusch, Tamara Müller, Dorothea Schulte, Janina Ehses, Michael Kiebler, Julia Hasler, et al. 'Choroid Plexus-derived MiR-204 Regulates the Number of Quiescent Neural Stem Cells in the Adult Brain'. *The EMBO Journal* 38, no. 17 (2 September 2019). https://doi.org/10.15252/embj.2018100481.

Li, Dong-Bin, Jing-Li Liu, Wei Wang, Xiu-Mei Luo, Xia Zhou, Jin-Pin Li, Xiao-Li Cao, Xiao-Hong Long, Jia-Gui Chen, and Chao Qin. 'Plasma Exosomal MiRNA-122-5p and MiR-300-3p as Potential Markers for Transient Ischaemic Attack in Rats'. *Frontiers in Aging Neuroscience* 10 (6 February 2018): 24. https://doi.org/10.3389/fnagi.2018.00024.

Li, Jian, Xiangnan Li, Xin Jiang, Mei Yang, Rui Yang, Geoffrey Burnstock, Zhenghua Xiang, and Hongbin Yuan. 'Microvesicles Shed from Microglia Activated by the P2X7-P38 Pathway Are Involved in Neuropathic Pain Induced by Spinal Nerve Ligation in Rats'. *Purinergic Signalling* 13, no. 1 (March 2017): 13–26. https://doi.org/10.1007/s11302-016-9537-0.

Li, Junjun, Hongliang Yuan, Hao Xu, Hongyang Zhao, and Nanxiang Xiong. 'Hypoxic Cancer-Secreted Exosomal MiR-182-5p Promotes Glioblastoma Angiogenesis by Targeting Kruppel-like Factor 2 and 4'. *Molecular Cancer Research* 18, no. 8 (1 August 2020): 1218–31. https://doi.org/10.1158/1541-7786.MCR-19-0725.

Li, Meng, Liu Huang, Joyce Chen, Fangfang Ni, Yating Zhang, and Fei Liu. 'Isolation of Exosome Nanoparticles from Human Cerebrospinal Fluid for Proteomic Analysis'. *ACS Applied Nano Materials* 4, no. 4 (23 April 2021): 3351–59. https://doi.org/10.1021/acsanm.0c02622.

Li, Yongang, Jiachen Gu, Youbing Mao, Xijia Wang, Zongshan Li, Xiaomin Xu, Huimin Chen, and Yaxing Gui. 'Cerebrospinal Fluid Extracellular Vesicles with Distinct Properties in Autoimmune Encephalitis and Herpes Simplex Encephalitis'. *Molecular Neurobiology* 59, no. 4 (April 2022): 2441–55. https://doi.org/10.1007/s12035-021-02705-2.

Li, Yuchen, Xigan He, Qin Li, Hongyan Lai, Hena Zhang, Zhixiang Hu, Yan Li, and Shenglin Huang. 'EV-Origin: Enumerating the Tissue-Cellular Origin of Circulating Extracellular Vesicles Using ExLR Profile'. *Computational and Structural Biotechnology Journal* 18 (2020): 2851–59. https://doi.org/10.1016/j.csbj.2020.10.002.

Lin, Yi-Wei, Jennifer Nhieu, Chin-Wen Wei, Yu-Lung Lin, Hiroyuki Kagechika, and Li-Na Wei. 'Regulation of Exosome Secretion by Cellular Retinoic Acid Binding Protein 1 Contributes to Systemic Anti-Inflammation'. *Cell Communication and Signaling* 19, no. 1 (December 2021): 69. https://doi.org/10.1186/s12964-021-00751-w.

Liu, Chen Geng, Chuang Meng, Ying Li, Yao Lu, Yue Zhao, and Pei Chang Wang. 'MicroRNA-135a in ABCA1-Labeled Exosome Is a Serum Biomarker Candidate for Alzheimer's Disease'. *Biomedical and Environmental Sciences* 34, no. 1 (2021): 19–28. https://doi.org/10.3967/bes2021.004.

Liu, Chen-Geng, Yue Zhao, Yao Lu, and Pei-Chang Wang. 'ABCA1-Labeled Exosomes in Serum Contain Higher MicroRNA-193b Levels in Alzheimer's Disease'. Edited by Dong-Yuan Cao. *BioMed Research International* 2021 (8 March 2021): 1–10. https://doi.org/10.1155/2021/5450397.

Liu, Xiaofeng, Zhengqing Zhao, Ruihua Ji, Jiao Zhu, Qian-Qian Sui, Gillian E. Knight, Geoffrey Burnstock, Cheng He, Hongbin Yuan, and Zhenghua Xiang. 'Inhibition of P2X7 Receptors Improves Outcomes after Traumatic Brain Injury in Rats'. *Purinergic Signalling* 13, no. 4 (December 2017): 529–44. https://doi.org/10.1007/s11302-017-9579-y.

Longobardi, Antonio, Roland Nicsanu, Sonia Bellini, Rosanna Squitti, Marcella Catania, Pietro Tiraboschi, Claudia Saraceno, et al. 'Cerebrospinal Fluid EV Concentration and Size Are Altered in Alzheimer's Disease and Dementia with Lewy Bodies'. *Cells* 11, no. 3 (28 January 2022): 462. https://doi.org/10.3390/cells11030462.

López-Pérez, Óscar, David Sanz-Rubio, Adelaida Hernaiz, Marina Betancor, Alicia Otero, Joaquín Castilla, Olivier Andréoletti, et al. 'Cerebrospinal Fluid and Plasma Small Extracellular Vesicles and MiRNAs as Biomarkers for Prion Diseases'. *International Journal of Molecular Sciences* 22, no. 13 (25 June 2021): 6822. https://doi.org/10.3390/ijms22136822.

Luo, XiuMei, Wei Wang, DongBin Li, Chen Xu, Bao Liao, FengMei Li, Xia Zhou, Wu Qin, and Jingli Liu. 'Plasma Exosomal MiR-450b-5p as a Possible Biomarker and Therapeutic Target for Transient Ischaemic Attacks in Rats'. *Journal of Molecular Neuroscience* 69, no. 4 (December 2019): 516–26. https://doi.org/10.1007/s12031-019-01341-9.

Madhankumar, A.B., Oliver D. Mrowczynski, Suhag R. Patel, Cody L. Weston, Brad E. Zacharia, Michael J. Glantz, Christopher A. Siedlecki, Li-Chong Xu, and James R. Connor. 'Interleukin-13 Conjugated Quantum Dots for Identification of Glioma Initiating Cells and Their Extracellular Vesicles'. *Acta Biomaterialia* 58 (August 2017): 205–13. https://doi.org/10.1016/j.actbio.2017.06.002.

Manek, Rachna, Ahmed Moghieb, Zhihui Yang, Dhwani Kumar, Firas Kobessiy, George Anis Sarkis, Vijaya Raghavan, and Kevin K.W. Wang. 'Protein Biomarkers and Neuroproteomics Characterization of Microvesicles/Exosomes from Human Cerebrospinal Fluid Following Traumatic Brain Injury'. *Molecular Neurobiology* 55, no. 7 (July 2018): 6112–28. https://doi.org/10.1007/s12035-017-0821-y.

Masvekar, Ruturaj, Jordan Mizrahi, John Park, Peter R. Williamson, and Bibiana Bielekova. 'Quantifications of CSF Apoptotic Bodies Do Not Provide Clinical Value in Multiple Sclerosis'. *Frontiers in Neurology* 10 (26 November 2019): 1241. https://doi.org/10.3389/fneur.2019.01241.

McKeever, Paul M., Raphael Schneider, Foad Taghdiri, Anna Weichert, Namita Multani, Robert A. Brown, Adam L. Boxer, et al. 'MicroRNA Expression Levels Are Altered in the Cerebrospinal Fluid of Patients with Young-Onset Alzheimer's Disease'. *Molecular Neurobiology* 55, no. 12 (December 2018): 8826–41. https://doi.org/10.1007/s12035-018-1032-x.

Minakaki, Georgia, Stefanie Menges, Agnes Kittel, Evangelia Emmanouilidou, Iris Schaeffner, Katalin Barkovits, Anna Bergmann, et al. 'Autophagy Inhibition Promotes SNCA/Alpha-Synuclein Release and Transfer via Extracellular Vesicles with a Hybrid Autophagosome-Exosome-like Phenotype'. *Autophagy* 14, no. 1 (2 January 2018): 98–119. https://doi.org/10.1080/15548627.2017.1395992.

Muraoka, Satoshi, Mark P. Jedrychowski, Harutsugu Tatebe, Annina M. DeLeo, Seiko Ikezu, Takahiko Tokuda, Steven P. Gygi, Robert A. Stern, and Tsuneya Ikezu. 'Proteomic Profiling of Extracellular Vesicles Isolated From Cerebrospinal Fluid of Former National Football League Players at Risk for Chronic Traumatic Encephalopathy'. *Frontiers in Neuroscience* 13 (9 October 2019): 1059. https://doi.org/10.3389/fnins.2019.01059.

Muraoka, Satoshi, Mark P. Jedrychowski, Kiran Yanamandra, Seiko Ikezu, Steven P. Gygi, and Tsuneya Ikezu. 'Proteomic Profiling of Extracellular Vesicles Derived from Cerebrospinal Fluid of Alzheimer's Disease Patients: A Pilot Study'. *Cells* 9, no. 9 (25 August 2020): 1959. https://doi.org/10.3390/cells9091959.

Norman, Maia, Dmitry Ter-Ovanesyan, Wendy Trieu, Roey Lazarovits, Emma J. K. Kowal, Ju Hyun Lee, Alice S. Chen-Plotkin, Aviv Regev, George M. Church, and David R. Walt. 'L1CAM Is Not Associated with Extracellular Vesicles in Human Cerebrospinal Fluid or Plasma'. *Nature Methods* 18, no. 6 (June 2021): 631–34. https://doi.org/10.1038/s41592-021-01174-8.

Otake, Kentaro, Hidenori Kamiguchi, and Yoshihiko Hirozane. 'Identification of Biomarkers for Amyotrophic Lateral Sclerosis by Comprehensive Analysis of Exosomal MRNAs in Human Cerebrospinal Fluid'. *BMC Medical Genomics* 12, no. 1 (December 2019): 7. https://doi.org/10.1186/s12920-019-0473-z.

Pieragostino, Damiana, Ilaria Cicalini, Paola Lanuti, Eva Ercolino, Maria di Ioia, Mirco Zucchelli, Romina Zappacosta, et al. 'Enhanced Release of Acid Sphingomyelinase-Enriched Exosomes Generates a Lipidomics Signature in CSF of Multiple Sclerosis Patients'. *Scientific Reports* 8, no. 1 (December 2018): 3071. https://doi.org/10.1038/s41598-018-21497-5.

Pieragostino, Damiana, Paola Lanuti, Ilaria Cicalini, Maria Concetta Cufaro, Fausta Ciccocioppo, Maurizio Ronci, Pasquale Simeone, et al. 'Proteomics Characterization of Extracellular Vesicles Sorted by Flow Cytometry Reveals a Disease-Specific Molecular Cross-Talk from Cerebrospinal Fluid and Tears in Multiple Sclerosis'. *Journal of Proteomics* 204 (July 2019): 103403. https://doi.org/10.1016/j.jprot.2019.103403.

Pisa, Marco, Tommaso Croese, Gloria Dalla Costa, Simone Guerrieri, Su-Chun Huang, Annamaria Finardi, Lorena Fabbella, et al. 'Subclinical Anterior Optic Pathway Involvement in Early Multiple Sclerosis and Clinically Isolated Syndromes'. *Brain* 144, no. 3 (12 April 2021): 848–62. https://doi.org/10.1093/brain/awaa458.

Prada, Ilaria, Martina Gabrielli, Elena Turola, Alessia Iorio, Giulia D'Arrigo, Roberta Parolisi, Mariacristina De Luca, et al. 'Glia-to-Neuron Transfer of MiRNAs via Extracellular Vesicles: A New Mechanism Underlying Inflammation-Induced Synaptic Alterations'. *Acta Neuropathologica* 135, no. 4 (April 2018): 529–50. https://doi.org/10.1007/s00401-017-1803-x.

Prieto-Fernández, Endika, Ana María Aransay, Félix Royo, Esperanza González, Juan José Lozano, Borja Santos-Zorrozua, Nuria Macias-Camara, et al. 'A Comprehensive Study of Vesicular and Non-Vesicular MiRNAs from a Volume of Cerebrospinal Fluid Compatible with Clinical Practice'. *Theranostics* 9, no. 16 (2019): 4567–79. https://doi.org/10.7150/thno.31502.

Qi, Yanhua, Chuandi Jin, Wei Qiu, Rongrong Zhao, Shaobo Wang, Boyan Li, Zongpu Zhang, et al. 'The Dual Role of Glioma Exosomal MicroRNAs: Glioma Eliminates Tumor Suppressor MiR-1298-5p via Exosomes to Promote Immunosuppressive Effects of MDSCs'. *Cell Death & Disease* 13, no. 5 (May 2022): 426. https://doi.org/10.1038/s41419-022-04872-z.

Qiu, Wei, Xiaofan Guo, Boyan Li, Jian Wang, Yanhua Qi, Zihang Chen, Rongrong Zhao, et al. 'Exosomal MiR-1246 from Glioma Patient Body Fluids Drives the Differentiation and Activation of Myeloid-Derived Suppressor Cells'. *Molecular Therapy* 29, no. 12 (December 2021): 3449–64. https://doi.org/10.1016/j.ymthe.2021.06.023.

Ragonese, Paolo, Italia Liegro, Gabriella Schiera, Giuseppe Salemi, Sabrina Realmuto, Carlo Liegro, and Patrizia Proia. 'Toxic Effects on Astrocytes of Extracellular Vesicles from CSF of Multiple Sclerosis Patients: A Pilot in Vitro Study'. *Polish Journal of Pathology* 71, no. 3 (2020): 270–76. https://doi.org/10.5114/pjp.2020.99794.

Raoof, Rana, Eva M. Jimenez-Mateos, Sebastian Bauer, Björn Tackenberg, Felix Rosenow, Johannes Lang, Müjgan Dogan Onugoren, et al. 'Cerebrospinal Fluid MicroRNAs Are Potential Biomarkers of Temporal Lobe Epilepsy and Status Epilepticus'. *Scientific Reports* 7, no. 1 (December 2017): 3328. https://doi.org/10.1038/s41598-017-02969-6.

Rather, Hilal A., Shalini Mishra, Yixin Su, Ashish Kumar, Sangeeta Singh, Biswapriya B. Misra, Jingyun Lee, et al. 'Mass Spectrometry-Based Proteome Profiling of Extracellular Vesicles Derived from the Cerebrospinal Fluid of Adult Rhesus Monkeys Exposed to Cocaine throughout Gestation'. *Biomolecules* 12, no. 4 (28 March 2022): 510. https://doi.org/10.3390/biom12040510.

Raval, Ami P., Camila C. Martinez, Nancy H. Mejias, and Juan Pablo de Rivero Vaccari. 'Sexual Dimorphism in Inflammasome-Containing Extracellular Vesicles and the Regulation of Innate Immunity in the Brain of Reproductive Senescent Females'. *Neurochemistry International* 127 (July 2019): 29–37. https://doi.org/10.1016/j.neuint.2018.11.018.

Riancho, Javier, José Luis Vázquez-Higuera, Ana Pozueta, Carmen Lage, Martha Kazimierczak, María Bravo, Miguel Calero, et al. 'MicroRNA Profile in Patients with Alzheimer's Disease: Analysis of MiR-9-5p and MiR-598 in Raw and Exosome Enriched Cerebrospinal Fluid Samples'. *Journal of Alzheimer's Disease* 57, no. 2 (21 March 2017): 483–91. https://doi.org/10.3233/JAD-161179.

Rider, Mark A., Stephanie N. Hurwitz, and David G. Meckes. 'ExtraPEG: A Polyethylene Glycol-Based Method for Enrichment of Extracellular Vesicles'. *Scientific Reports* 6, no. 1 (July 2016): 23978. https://doi.org/10.1038/srep23978.

Sanchez, Isabella I., Thai B. Nguyen, Whitney E. England, Ryan G. Lim, Anthony Q. Vu, Ricardo Miramontes, Lauren M. Byrne, et al. 'Huntington's Disease Mice and Human Brain Tissue Exhibit Increased G3BP1 Granules and TDP43 Mislocalization'. *Journal of Clinical Investigation* 131, no. 12 (15 June 2021): e140723. https://doi.org/10.1172/JCI140723.

Sandau, Ursula S., Trevor J. McFarland, Sierra J. Smith, Douglas R. Galasko, Joseph F. Quinn, and Julie A. Saugstad. 'Differential Effects of APOE Genotype on MicroRNA Cargo of Cerebrospinal Fluid Extracellular Vesicles in Females With Alzheimer's Disease Compared to Males'. *Frontiers in Cell and Developmental Biology* 10 (27 April 2022): 864022. https://doi.org/10.3389/fcell.2022.864022.

Saugstad, Julie A., Theresa A. Lusardi, Kendall R. Van Keuren-Jensen, Jay I. Phillips, Babett Lind, Christina A. Harrington, Trevor J. McFarland, et al. 'Analysis of Extracellular RNA in Cerebrospinal Fluid'. *Journal of Extracellular Vesicles* 6, no. 1 (1 December 2017): 1317577. https://doi.org/10.1080/20013078.2017.1317577.

Schneider, Raphael, Paul McKeever, TaeHyung Kim, Caroline Graff, John Cornelis van Swieten, Anna Karydas, Adam Boxer, et al. 'Downregulation of Exosomal MiR-204-5p and MiR-632 as a Biomarker for FTD: A GENFI Study'. *Journal of Neurology, Neurosurgery & Psychiatry* 89, no. 8 (August 2018): 851–58. https://doi.org/10.1136/jnnp-2017-317492.

Shi, Rui, Pei-Yin Wang, Xin-Yi Li, Jian-Xin Chen, Yan Li, Xin-Zhong Zhang, Chen-Guang Zhang, et al. 'Exosomal Levels of MiRNA-21 from Cerebrospinal Fluids Associated with Poor Prognosis and Tumor Recurrence of Glioma Patients'. *Oncotarget* 6, no. 29 (29 September 2015): 26971–81. https://doi.org/10.18632/oncotarget.4699.

Sjoqvist, Sebastian, and Kentaro Otake. 'A Pilot Study Using Proximity Extension Assay of Cerebrospinal Fluid and Its Extracellular Vesicles Identifies Novel Amyotrophic Lateral Sclerosis Biomarker Candidates'. *Biochemical and Biophysical Research Communications* 613 (July 2022): 166–73. https://doi.org/10.1016/j.bbrc.2022.04.127.

Sjoqvist, Sebastian, Kentaro Otake, and Yoshihiko Hirozane. 'Analysis of Cerebrospinal Fluid Extracellular Vesicles by Proximity Extension Assay: A Comparative Study of Four Isolation Kits'. *International Journal of Molecular Sciences* 21, no. 24 (10 December 2020): 9425. https://doi.org/10.3390/ijms21249425.

Skalnikova, Bohuslavova, Turnovcova, Juhasova, Juhas, Rodinova, and Vodicka. ‘Isolation and Characterization of Small Extracellular Vesicles from Porcine Blood Plasma, Cerebrospinal Fluid, and Seminal Plasma’. *Proteomes* 7, no. 2 (25 April 2019): 17. https://doi.org/10.3390/proteomes7020017.

Soares Martins, Tânia, José Catita, Ilka Martins Rosa, Odete A. B. da Cruz e Silva, and Ana Gabriela Henriques. ‘Exosome Isolation from Distinct Biofluids Using Precipitation and Column-Based Approaches’. Edited by Guo-Chang Fan. *PLOS ONE* 13, no. 6 (11 June 2018): e0198820. https://doi.org/10.1371/journal.pone.0198820.

Spaull, Robert, Bryony McPherson, Andriana Gialeli, Aled Clayton, James Uney, Axel Heep, and Óscar Cordero-Llana. ‘Exosomes Populate the Cerebrospinal Fluid of Preterm Infants with Post-haemorrhagic Hydrocephalus’. *International Journal of Developmental Neuroscience* 73, no. 1 (April 2019): 59–65. https://doi.org/10.1016/j.ijdevneu.2019.01.004.

Spitzer, Philipp, Linda-Marie Mulzer, Timo Jan Oberstein, Luis Enrique Munoz, Piotr Lewczuk, Johannes Kornhuber, Martin Herrmann, and Juan Manuel Maler. ‘Microvesicles from Cerebrospinal Fluid of Patients with Alzheimer’s Disease Display Reduced Concentrations of Tau and APP Protein’. *Scientific Reports* 9, no. 1 (December 2019): 7089. https://doi.org/10.1038/s41598-019-43607-7.

Stuendl, Anne, Marcel Kunadt, Niels Kruse, Claudia Bartels, Wiebke Moebius, Karin M. Danzer, Brit Mollenhauer, and Anja Schneider. ‘Induction of α-Synuclein Aggregate Formation by CSF Exosomes from Patients with Parkinson’s Disease and Dementia with Lewy Bodies’. *Brain* 139, no. 2 (February 2016): 481–94. https://doi.org/10.1093/brain/awv346.

Tan, Ning, Shuiwang Hu, Zhen Hu, Zhouli Wu, and Bin Wang. ‘Quantitative Proteomic Characterization of Microvesicles/Exosomes from the Cerebrospinal Fluid of Patients with Acute Bilirubin Encephalopathy’. *Molecular Medicine Reports* 22, no. 2 (28 May 2020): 1257–68. https://doi.org/10.3892/mmr.2020.11194.

Tan, Yi Jayne, Benjamin Y.X. Wong, Ramanathan Vaidyanathan, Sivaramapanicker Sreejith, Sook Yoong Chia, Nagaendran Kandiah, Adeline S.L. Ng, and Li Zeng. ‘Altered Cerebrospinal Fluid Exosomal MicroRNA Levels in Young-Onset Alzheimer’s Disease and Frontotemporal Dementia’. *Journal of Alzheimer’s Disease Reports* 5, no. 1 (28 October 2021): 805–13. https://doi.org/10.3233/ADR-210311.

Ter-Ovanesyan, Dmitry, Maia Norman, Roey Lazarovits, Wendy Trieu, Ju-Hyun Lee, George M Church, and David R Walt. ‘Framework for Rapid Comparison of Extracellular Vesicle Isolation Methods’. *ELife* 10 (16 November 2021): e70725. https://doi.org/10.7554/eLife.70725.

Thakur, Abhimanyu, Chen Xu, Wing Kar Li, Guangyu Qiu, Bing He, Siu-Pang Ng, Chi-Man Lawrence Wu, and Youngjin Lee. ‘In Vivo Liquid Biopsy for Glioblastoma Malignancy by the AFM and LSPR Based Sensing of Exosomal CD44 and CD133 in a Mouse Model’. *Biosensors and Bioelectronics* 191 (November 2021): 113476. https://doi.org/10.1016/j.bios.2021.113476.

Thompson, Alexander G., Elizabeth Gray, Imre Mager, Roman Fischer, Marie-Laëtitia Thézénas, Philip D Charles, Kevin Talbot, et al. ‘UFLC-Derived CSF Extracellular Vesicle Origin and Proteome’. *PROTEOMICS*, 10 December 2018, 1800257. https://doi.org/10.1002/pmic.201800257.

Thompson, Alexander G., Elizabeth Gray, Imre Mäger, Marie-Laëtitia Thézénas, Philip D. Charles, Kevin Talbot, Roman Fischer, Benedikt M. Kessler, Mathew Wood, and Martin R. Turner. ‘CSF Extracellular Vesicle Proteomics Demonstrates Altered Protein Homeostasis in Amyotrophic Lateral Sclerosis’. *Clinical Proteomics* 17, no. 1 (December 2020): 31. https://doi.org/10.1186/s12014-020-09294-7.

Tsutsui, Taishi, Hironori Kawahara, Ryouken Kimura, Yu Dong, Shabierjiang Jiapaer, Hemragul Sabit, Jiakang Zhang, Takeshi Yoshida, Mitsutoshi Nakada, and Rikinari Hanayama. 'Glioma-Derived Extracellular Vesicles Promote Tumor Progression by Conveying WT1'. *Carcinogenesis* 41, no. 9 (24 September 2020): 1238–45. https://doi.org/10.1093/carcin/bgaa052.

Utz, Janine, Judith Berner, Luis Enrique Muñoz, Timo Jan Oberstein, Johannes Kornhuber, Martin Herrmann, Juan Manuel Maler, and Philipp Spitzer. 'Cerebrospinal Fluid of Patients With Alzheimer's Disease Contains Increased Percentages of Synaptophysin-Bearing Microvesicles'. *Frontiers in Aging Neuroscience* 13 (6 July 2021): 682115. https://doi.org/10.3389/fnagi.2021.682115.

Vacchi, Elena, Jacopo Burrello, Alessio Burrello, Sara Bolis, Silvia Monticone, Lucio Barile, Alain Kaelin-Lang, and Giorgia Melli. 'Profiling Inflammatory Extracellular Vesicles in Plasma and Cerebrospinal Fluid: An Optimized Diagnostic Model for Parkinson's Disease'. *Biomedicines* 9, no. 3 (25 February 2021): 230. https://doi.org/10.3390/biomedicines9030230.

Van Hoecke, Lien, Caroline Van Cauwenberghe, Kristina Dominko, Griet Van Imschoot, Elien Van Wonterghem, Jonas Castelein, Junhua Xie, et al. 'Involvement of the Choroid Plexus in the Pathogenesis of Niemann-Pick Disease Type C'. *Frontiers in Cellular Neuroscience* 15 (15 October 2021): 757482. https://doi.org/10.3389/fncel.2021.757482.

Vandendriessche, Charysse, Sriram Balusu, Caroline Van Cauwenberghe, Marjana Brkic, Marie Pauwels, Nele Plehiers, Arnout Bruggeman, et al. 'Importance of Extracellular Vesicle Secretion at the Blood–Cerebrospinal Fluid Interface in the Pathogenesis of Alzheimer's Disease'. *Acta Neuropathologica Communications* 9, no. 1 (December 2021): 143. https://doi.org/10.1186/s40478-021-01245-z.

Wang, Gang, Yunyu Wen, Oluwasijibomi Damola Faleti, Qingshun Zhao, Jingping Liu, Guozhong Zhang, Mingzhou Li, Songtao Qi, Wenfeng Feng, and Xiaoming Lyu. 'A Panel of Exosome-Derived MiRNAs of Cerebrospinal Fluid for the Diagnosis of Moyamoya Disease'. *Frontiers in Neuroscience* 14 (25 September 2020): 548278. https://doi.org/10.3389/fnins.2020.548278.

Wang, Ming, Yang Cai, Yong Peng, Bo Xu, Wentao Hui, and Yugang Jiang. 'Exosomal LGALS9 in the Cerebrospinal Fluid of Glioblastoma Patients Suppressed Dendritic Cell Antigen Presentation and Cytotoxic T-Cell Immunity'. *Cell Death & Disease* 11, no. 10 (October 2020): 896. https://doi.org/10.1038/s41419-020-03042-3.

Wang, Shijie, Kaela Kelly, Jonathan M. Brotchie, James B. Koprich, and Andrew B. West. 'Exosome Markers of LRRK2 Kinase Inhibition'. *Npj Parkinson's Disease* 6, no. 1 (December 2020): 32. https://doi.org/10.1038/s41531-020-00138-7.

Wang, Shijie, Zhiyong Liu, Tao Ye, Omar S. Mabrouk, Tyler Maltbie, Jan Aasly, and Andrew B. West. 'Elevated LRRK2 Autophosphorylation in Brain-Derived and Peripheral Exosomes in LRRK2 Mutation Carriers'. *Acta Neuropathologica Communications* 5, no. 1 (December 2017): 86. https://doi.org/10.1186/s40478-017-0492-y.

Wang, Yipeng, Varun Balaji, Senthilvelrajan Kaniyappan, Lars Krüger, Stephan Irsen, Katharina Tepper, RamReddy Chandupatla, et al. 'The Release and Trans-Synaptic Transmission of Tau via Exosomes'. *Molecular Neurodegeneration* 12, no. 1 (December 2017): 5. https://doi.org/10.1186/s13024-016-0143-y.

Wei, Zhiyun, Arsen O. Batagov, Sergio Schinelli, Jintu Wang, Yang Wang, Rachid El Fatimy, Rosalia Rabinovsky, et al. 'Coding and Noncoding Landscape of Extracellular RNA Released by Human Glioma Stem Cells'. *Nature Communications* 8, no. 1 (December 2017): 1145. https://doi.org/10.1038/s41467-017-01196-x.

Welton, Joanne L., Samantha Loveless, Timothy Stone, Chris von Ruhland, Neil P. Robertson, and Aled Clayton. 'Cerebrospinal Fluid Extracellular Vesicle Enrichment for Protein Biomarker Discovery in Neurological Disease; Multiple Sclerosis'. *Journal of Extracellular Vesicles* 6, no. 1 (1 December 2017): 1369805. https://doi.org/10.1080/20013078.2017.1369805.

Wilson, M. E., C. L. Holz, A. K. Kopec, J. J. Dau, J. P. Luyendyk, and G. Soboll Hussey. 'Coagulation Parameters Following Equine Herpesvirus Type 1 Infection in Horses'. *Equine Veterinary Journal* 51, no. 1 (January 2019): 102–7. https://doi.org/10.1111/evj.12843.

Xu, Hao, Ming Li, Ziwen Pan, Zongpu Zhang, Zijie Gao, Rongrong Zhao, Boyan Li, et al. 'MiR-3184-3p Enriched in Cerebrospinal Fluid Exosomes Contributes to Progression of Glioma and Promotes M2-like Macrophage Polarization'. *Cancer Science*, 4 May 2022, cas.15372. https://doi.org/10.1111/cas.15372.

Yagi, Yohsuke, Takuya Ohkubo, Hideya Kawaji, Akira Machida, Haruka Miyata, Saori Goda, Sugata Roy, Yoshihide Hayashizaki, Harukazu Suzuki, and Takanori Yokota. 'Next-Generation Sequencing-Based Small RNA Profiling of Cerebrospinal Fluid Exosomes'. *Neuroscience Letters* 636 (January 2017): 48–57. https://doi.org/10.1016/j.neulet.2016.10.042.

Yang, Yue, C. Dirk Keene, Elaine R. Peskind, Douglas R. Galasko, Shu-Ching Hu, Eiron Cudaback, Angela M. Wilson, et al. 'Cerebrospinal Fluid Particles in Alzheimer Disease and Parkinson Disease'. *Journal of Neuropathology & Experimental Neurology* 74, no. 7 (July 2015): 672–87. https://doi.org/10.1097/NEN.0000000000000207.

Yao, Yang, Xinggen Fang, Jinlong Yuan, Feiyun Qin, Tao Yu, Dayong Xia, Zhenbao Li, and Niansheng Lai. 'Interleukin-6 in Cerebrospinal Fluid Small Extracellular Vesicles as a Potential Biomarker for Prognosis of Aneurysmal Subarachnoid Haemorrhage'. *Neuropsychiatric Disease and Treatment* Volume 17 (May 2021): 1423–31. https://doi.org/10.2147/NDT.S304394.

Yelick, Julia, Yuqin Men, Shijie Jin, Sabrina Seo, Francisco Espejo-Porras, and Yongjie Yang. 'Elevated Exosomal Secretion of MiR-124-3p from Spinal Neurons Positively Associates with Disease Severity in ALS'. *Experimental Neurology* 333 (November 2020): 113414. https://doi.org/10.1016/j.expneurol.2020.113414.
