## Supplementary figures for "Towards optimised extracellular vesicle proteomics from cerebrospinal fluid"

*Shared last author

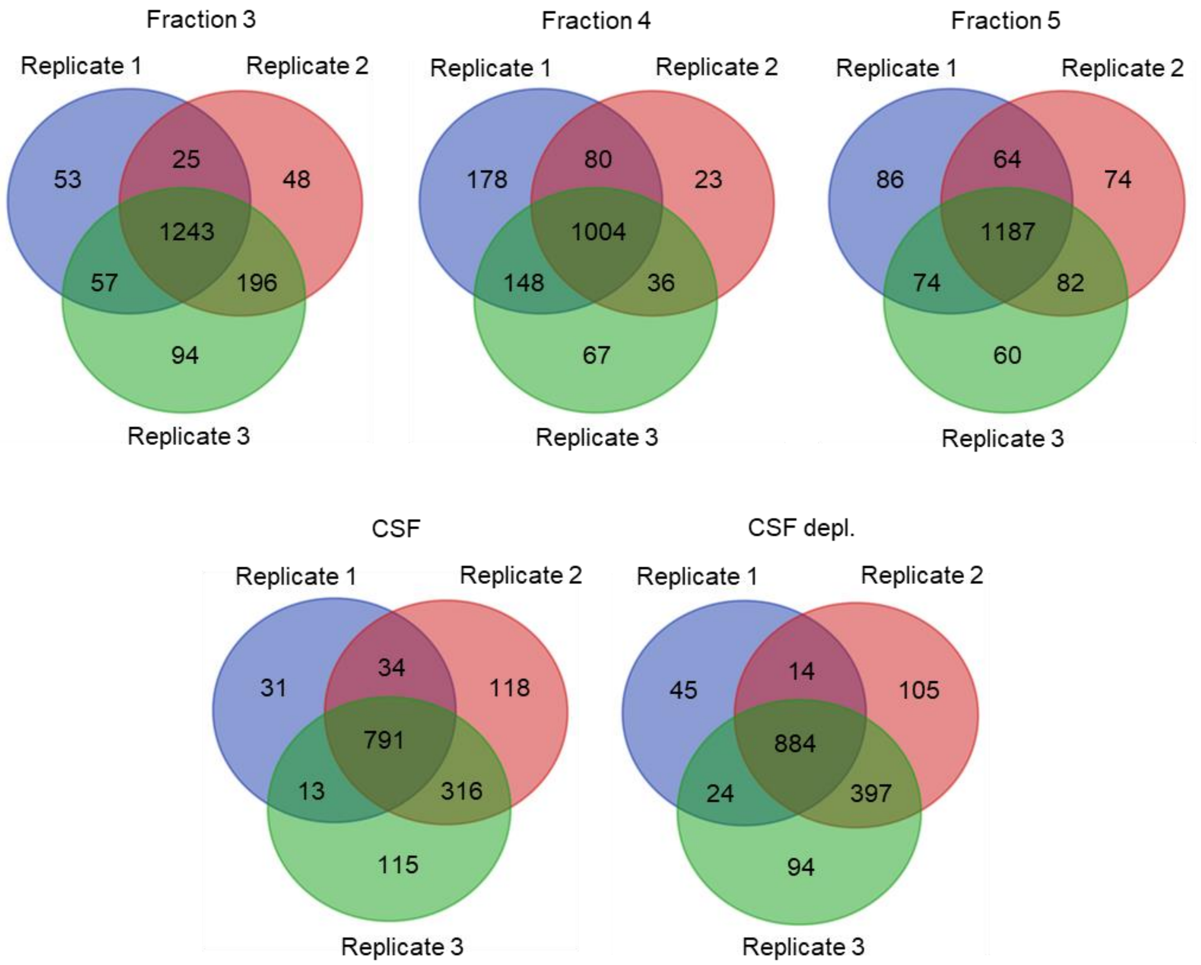


**Supplementary figure 1:** Venn diagrams of the identified proteins for each replicate of fractions 3-5 as well as CSF and CSF depl. Protein identification was done with 'matches between runs' active in MaxQuant. Abbreviations: CSF = cerebrospinal fluid, CSF depl. = CSF with depletion of highest abundant plasma proteins

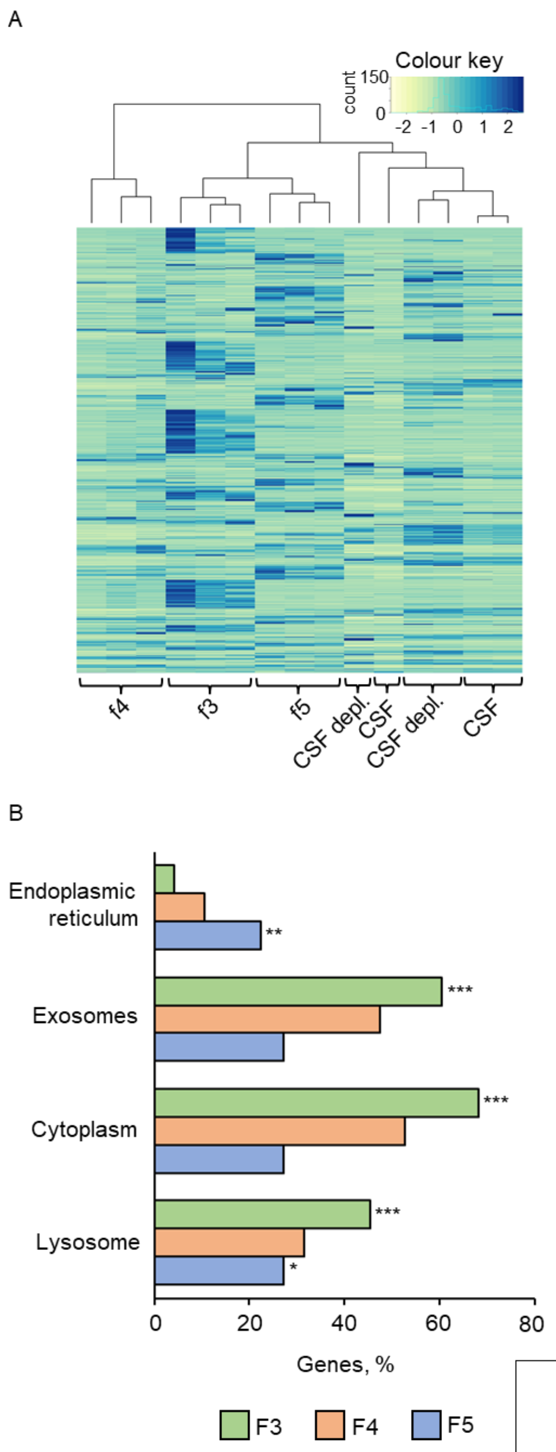


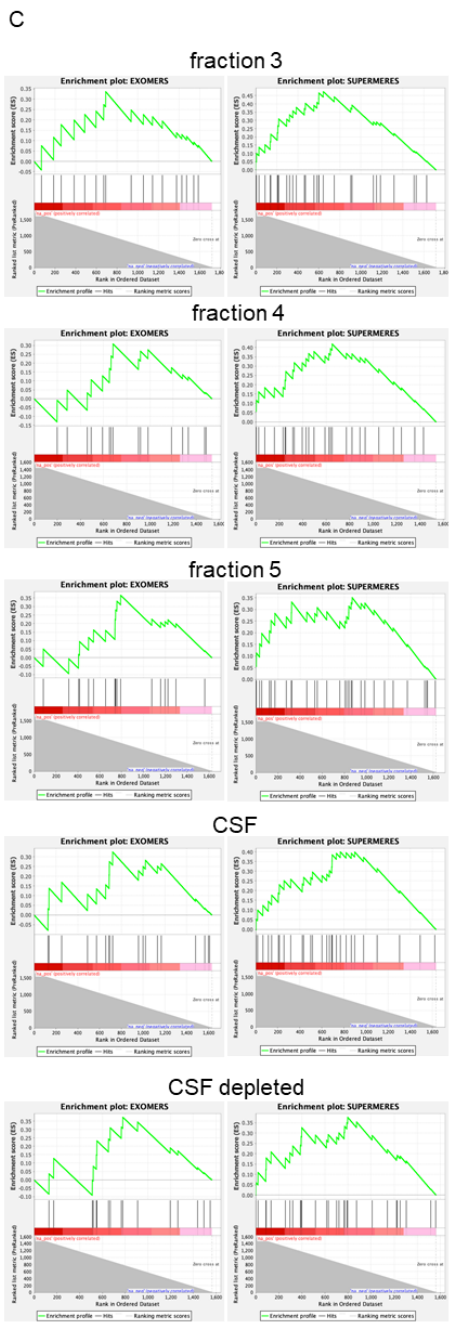


| | exomers | | | supermeres | | |
|---|---|---|---|---|---|---|
| | ES | NOM p-value | FDR q-val | ES | NOM p-value | FDR q-val |
| fraction 3 | 0.34 | 0.540 | 0.561 | 0.47 | 0.011 | 0.019 |
| fraction 4 | 0.31 | 0.700 | 0.746 | 0.42 | 0.077 | 0.108 |
| fraction 5 | 0.37 | 0.399 | 0.581 | 0.35 | 0.288 | 0.816 |
| csf | 0.32 | 0.571 | 0.598 | 0.40 | 0.112 | 0.327 |
| csf depl. | 0.37 | 0.396 | 0.580 | 0.38 | 0.231 | 0.612 |

**Supplementary figure 2: A.** Heatmap with clustering of all the proteins found in the samples. **B.** Pathway analysis of chosen cellular components of SEC fractions 3-5 using FunRich. **C.** GSEA of protein list for exomers and supermeres. Abbreviations: SEC = size-exclusion chromatography, GSEA = gene set enrichment analysis, CSF = cerebrospinal fluid, CSF depl. = CSF with depletion of highest abundant plasma proteins. *p≤0.05, **p≤0.01, ***p≤0.001.

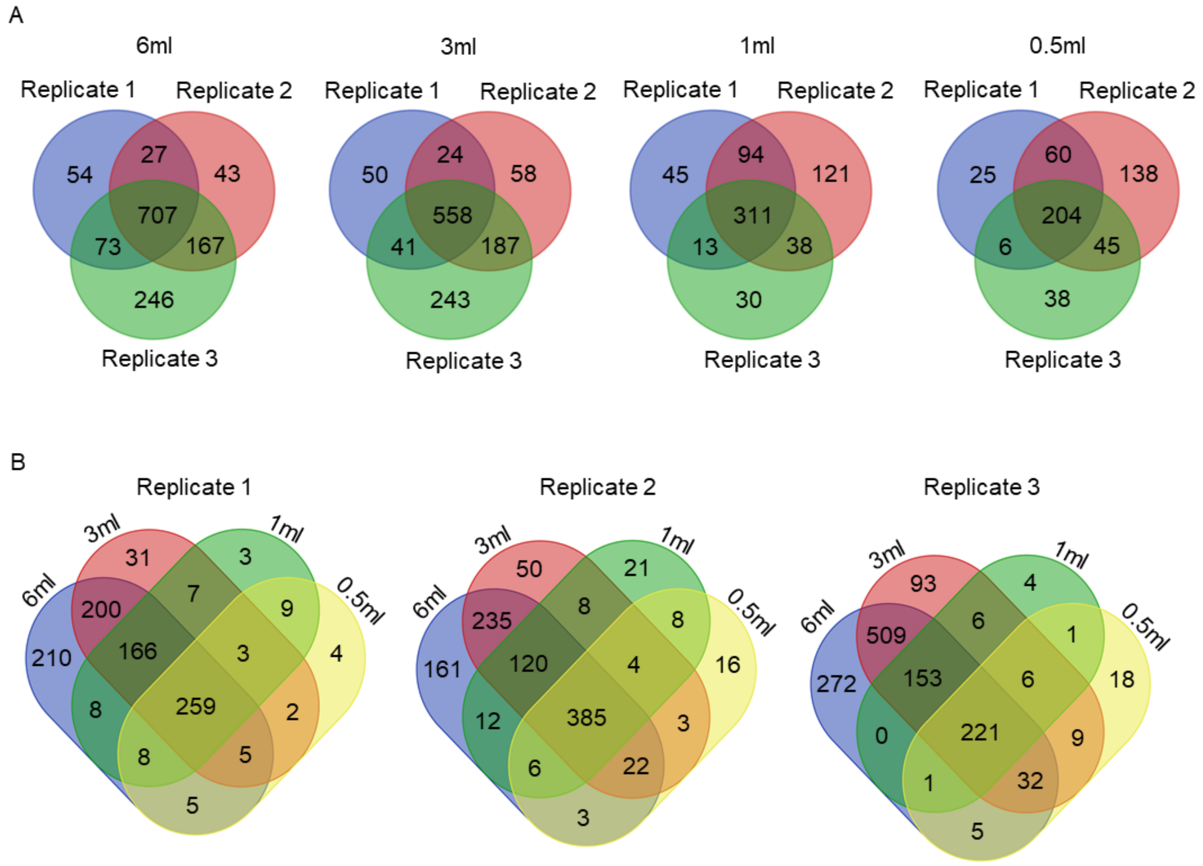


**Supplementary figure 3: A.** Shared proteins for the different starting volumes in each proteomics replicate. **B.** Shows the shared proteins between the different starting volumes in each proteomics replicate. Identification was done without the 'matches between runs' active in MaxQuant.
